# Neural Traces of Forgotten Memories Persist in Humans and are Behaviorally Relevant

**DOI:** 10.1101/2025.06.02.656652

**Authors:** Tom Willems, Konstantinos Zervas, Luzius Brogli, Finn Rabe, Andrea Federspiel, Katharina Henke

## Abstract

For a long time, forgetting has been taken as the dissipation of the neural memory traces (engrams). However, recent engram research in mice suggests that the engrams of forgotten memories do persist. This raises the question of whether engrams underlying human episodic memories also persist despite forgetting? And do forgotten memories influence human behavior implicitly? To address these questions, we used high-resolution functional magnetic resonance imaging at 7 Tesla to map the fate of 96 associative memories at the systems level from learning to a 30-minute and onward to a 24-hour memory test. Upon each retrieval attempt, participants indicated whether they remembered or forgot the memory. Univariate and multivariate analyses of the functional brain data revealed that the engrams of forgotten memories remain implemented in the episodic memory network and continue to influence the accuracy of guessing responses at the memory test. Overnight, the engrams of forgotten memories became implemented more deeply within bilateral hippocampus, while consciously accessible memories were neocorticalized overnight. The engrams of both consciously accessible and inaccessible (forgotten) memories shifted from the 30-minute to the 24-hour memory test within the right hippocampus and anterior cingulate gyrus as evidenced by the occurrence of pattern dissimilarities that supported correct retrieval responses at 24 hours. Hence, forgotten human episodic memories remain implemented in the episodic memory system and continue to influence decisions.

## INTRODUCTION

Most neuroscientific research focuses on learning and remembering, while the nature of forgetting has been largely overlooked. Here, we define forgetting as the insight that the retrieval of a previously accessible memory has failed (1–7). Indeed, healthy, adult participants are very sensitive to retrieval failures(*1*, *2*). Here, we investigate whether forgetting in episodic memory is merely a failure of conscious access to a memory or reflects the loss of the neural trace of a memory. Learning involves the strengthening of connections between coactivated neurons that are part of a neuronal ensemble (memory trace or engram), which is associated with a particular memory (*3*). Memories are encoded in sparse neural ensembles distributed across the entire brain (*4*). Following learning, engram cells that were active during learning are reactivated, which supports the strengthening (consolidation) of the memory. Forgetting is the flip side of memory consolidation. A common view is that forgetting is due to the dissipation of the entire engram due to neurogenesis, interference, and due to the brain’s tendency to degrade molecular and cellular memory traces leading to the disconnection of engram cells from the engram circuit (*5*). Recent engram research in rodents, however, suggests that it is unlikely that the engram of forgotten memories vanishes completely (*4*). Instead, forgetting may occur due to a reduced accessibility of engrams that silently persist.

Engram accessibility may depend on the intactness of the engram itself, on the systemic state of the individual, and on the quality of the retrieval cues (*6*). For example, Guskjolen and colleagues (*7*) found in young mice with natural infantile amnesia that optogenetic stimulation of hippocampal neurons tagged during contextual fear learning could recover forgotten memories for up to 3 months. Also, Yates et al. (*8*) found that human infantile amnesia is not a failure of encoding, but rather a failure of conscious retrieval of the memories that turn inaccessible due to post-encoding processes. Bolsius et al. (*9*) found that sleep-deprivation-induced amnesia in mice could be reversed by both optogenetic stimulation and the drug roflumilast, which acts by preventing the breakdown of enzymes involved in long-term potentiation. In adult mice, Autore and colleagues (*10*) showed that although natural retroactive interference during learning resulted in forgetting and decreased engram cell reactivation in the retrieval situation, optogenetic stimulation of previously labeled engram cells enabled memory expression. These discoveries beg the question of whether hippocampal and neocortical engram components underlying adult, human episodic memories would also persist despite forgetting and – if so – whether the forgotten memories associated with a persistent engram would continue to influence human behavior implicitly. Here, we tested these hypotheses making use of anterograde and retrograde interference during learning to increase forgetting rates in healthy young men and women. Indeed, much forgetting is likely due to interference – i.e., multiple memories encoded closely in time creating competition during consolidation (*5*, *11*, *12*). But why should the traces of forgotten memories remain implemented in the brain? Perhaps because the forgotten memories may regain importance for behavioral adjustment and survival when anticipating or being confronted with a similar situation later in life.

On a cellular level, the degree of reinstatement is dependent on the strength of connectivity between engram cells (*13*). Synaptic consolidation provides for an active strengthening of the connectivity between engram neurons within an engram component via cellular mechanisms like long-term-potentiation (*14*, *15*). Systems consolidation describes a time-dependent change of episodic memory representation. These changes are characterized by a strengthening of neocortical, mostly medial prefrontal, engram components (*16*, *17*). Over time, retrieval tends to (but not always does) become independent from the hippocampal engram component (*18–20*). Because engrams undergo changes at multiple levels of neural organization from synapses to cells to neuronal ensembles, it is possible to study engrams at the systems level in humans with high-resolution functional magnetic resonance imaging (fMRI) and multivariate analyses (*21*). The degree of engram reinstatement within human neocortex and hippocampus has been found to predict retrieval success and the amount of information retrieved (*22–24*). Besides reinstatement-related similarity, memory accuracy might also be linked to a transformation of the trace reflected by distinct activation patterns within the hippocampus and dissimilarity between the learning and retrieval events (*25–27*). This dissimilarity of learning- and retrieval-related activation patterns may be caused by the rapid transformation of cell-ensemble dynamics in the hippocampus (*28–30*), by the reconsolidation of memories following a reactivation (*31*), and by pattern separation processes that provide for an orthogonal representation of similar memories to prevent interference (*27*, *32*).

Learning- and retrieval-related activation patterns that underlie forgotten rather than remembered adult human episodic memories have not yet been examined to our knowledge. The reason for this neglect might be conceptual and definitional in nature: If forgetting from episodic memory were associated with persisting engrams that continue to influence behavior implicitly and are located in the same neural circuits that underlie consciously accessible memories, this would entail that episodic retrieval is not all (conscious) or none (absent). Yet, unconscious episodic memory retrieval does not exist in the framework of traditional models of human memory because these models link episodic memory to consciousness (*33–35*). Note that the literature on episodic-autobiographical memory retrieval describes direct and involuntary forms of autobiographical memory retrieval, which can occur without conscious effort and may in fact be elicited by unconscious processes (*36–38*). However, the act of retrieval can always be reported, meaning it is conscious. This links both direct and involuntary autobiographical memory retrieval to consciousness, in line with traditional models of human memory. Accordingly, forgetting/remembering is all or none. Newer views of episodic memory allow for unconscious episodic memory formation and retrieval (*39–41*). These views open the field conceptually to studying all shades of forgetting from losing conscious access to finally also losing unconscious access to an episodic memory. The finding of persistent neural engrams of once consciously acquired episodic memories, which subsequently lost conscious accessibility, would therefore inform traditional models of human episodic memory.

There is some animal and human evidence indicating that memory traces of forgotten episodic memories might indeed influence behavior implicitly. For example, mice that had learned a Morris water maze as infants forget the location of the platform as they grow. Nevertheless, they are significantly quicker at relearning the location of the platform than naive adult mice (*42*). Furthermore, previously learned and then forgotten images were still retrieved implicitly at 24 hours following learning and yielded fMRI-measured brain activation changes in regions that partly overlapped with regions activated during successful conscious image retrieval (*43*). In addition, there is substantial evidence for the feasibility of an unconscious retrieval of subliminally encoded and sleep-encoded associative memories in humans (*44–48*), bolstering the claim that episodic learning and retrieval can proceed outside of conscious awareness. In the current experiment, we therefore expected to observe behavioral evidence of episodic memory expression even when participants indicated that they had forgotten the memory and therefore needed to guess (i.e., above-chance guessing).

In this translational endeavor, we measured blood-oxygenation-level dependent (BOLD) brain activity in 40 healthy young adults with high-resolution fMRI at 7 Tesla. The increased signal-to-noise ratio of 7 Tesla fMRI improves the spatial resolution, which enhances the measurement of the (dis)similarity of activity patterns for individual memory traces over time (*21*). The large number of associations, which needed to be learned rapidly, was intended to maximize forgetting by anterograde and retrograde interference (*11*, *12*, *49*, *50*). Participants were imaged during learning, an immediate retrieval, a 30-minute retrieval, and a 24-hour retrieval. Most forgetting from episodic memory occurs during the first 24 hours following learning (*51–53*). Half of participants (N = 20) were examined with a small field-of-view (FOV) fMRI sequence that focused on the bilateral medial temporal lobe (MTL; 0.8 mm isotropic voxel size) to target the hippocampal engram components, and other half (N = 20) with a whole-brain fMRI sequence (1.1 mm isotropic voxel size) that targeted neocortical engram components. Engram components were revealed using univariate and multivariate (representational similarity) fMRI analyses. Remembered and forgotten memories were probed separately and in conjunction to each other.

While learning, we presented a series of 96 face-object combinations, each combination being shown only once for one-shot learning (Figure 1). The presented objects were sampled from two categories, organic (e.g., kiwi) and inorganic (e.g., stapler). Memories were prompted with two retrievals: Cued by the image of a previously presented face, participants indicated whether the face-associated object was organic or inorganic (category retrieval) and cued by the same face plus the original object and a foil object, participants chose the object that was originally presented with the face (2-alternative forced-choice recognition). The foil object had been presented with another face during learning. Category retrieval followed the learning of a face-object combination immediately to ensure that learning was successful, and it was repeated 30 minutes and 24 hours following learning to track forgetting. The recognition task was only given at 24 hours after learning and it always followed category retrieval. Following each retrieval/recognition response, participants indicated whether their just given response was 1) a pure guess because they had forgotten the memory, 2) an unsure response because the memory was neither fully forgotten nor fully accessible, or 3) a sure response because the memory was consciously accessible. We evaluated only the extremes, i.e., the guess responses (forgotten) and the sure responses (consciously accessible) comparing their underlying brain activations.

**Figure 1.**
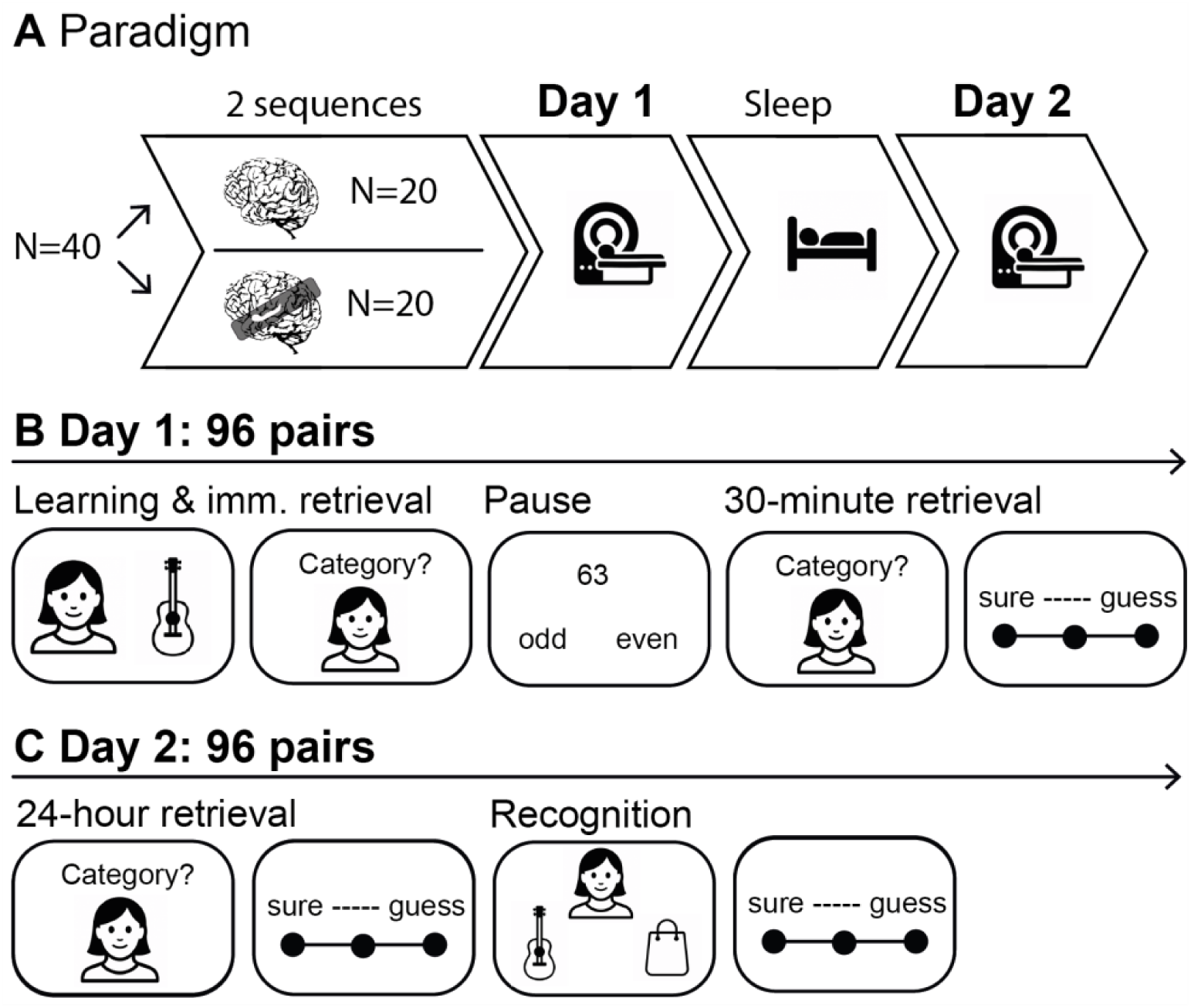
Experimental design. A) Overall procedure. Forty participants were randomly assigned to one of the two functional magnetic resonance imaging (fMRI) sequences, one fMRI sequence covering the whole brain and the other covering only the medial temporal lobe. Participants took part in two (f)MRI sessions, separated by 24 hours including one night of sleep. B) Day 1 procedure. Participants learned naturalistic face-object pairs (shown here as pictograms for illustration) and immediately retrieved the pairs with the category retrieval (imm. retrieval: immediate category retrieval) to assess whether successful learning occurred. Next, participants had a short pause of 5 minutes, during which they remained in the scanner and performed an odd-even task preventing them from actively rehearsing the face-object pairs. Following the short pause, participants took the 30-minute category retrieval. They chose the face-associated object category (organic or inorganic) and rated the confidence of their response (sure, unsure, guess). Due to the short pause and the pseudo-random shuffling of the face-object pairs, the interval between the one-trial learning and the 30-minute category retrieval was constant at 30 minutes for each pair. C) Day 2 procedure. Repetition of the category retrieval at 24 hours including the confidence rating, followed by an object recognition task and another confidence rating.

## RESULTS

### Behavioral results

Because the sample consisted of young and healthy men and women, there were almost no false memories in the sense of incorrect sure responses. On the immediate category retrieval, which was done to ensure correct memory formation, participants retrieved the object categories with a mean accuracy of 98% (SD = 2%). All responses were rated ‘sure’ (consciously retrieved). Those few trials that yielded an incorrect response were excluded from the subsequent analyses. Behavioral data of all 40 participants are summarized in Figure 2 and Table S1.

**Figure 2.**
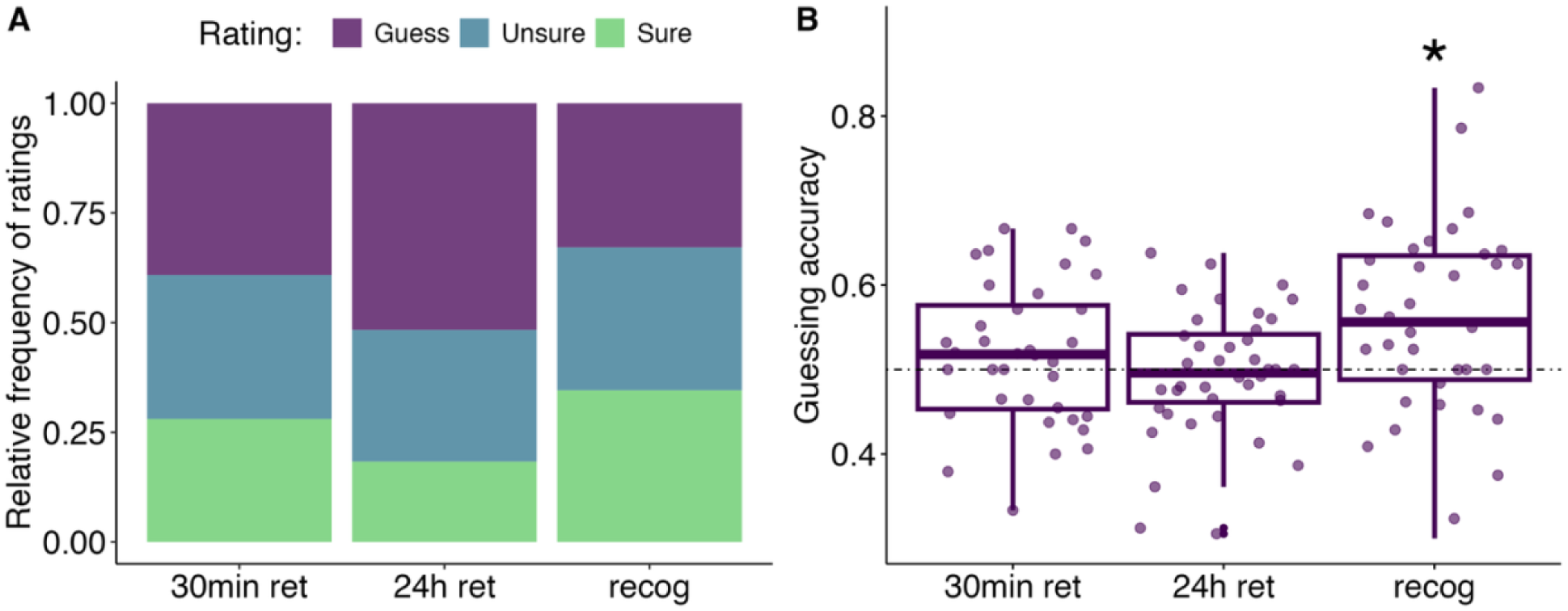
Behavioral results. A) Relative frequencies (bar sections) of confidence judgements across retrievals. Frequency of sure responses drops at the 24-hour category retrieval and rises again during recognition. Frequency of guess responses shows the opposite pattern of change. Frequency of unsure responses drops at the 24-hour category retrieval and remains constant between the 24-hour category retrieval and recognition. B) Guessing accuracy (percentage correct of all guess responses) on the category retrievals and the recognition. Above-chance accuracy was observed only for guess responses given on the recognition task (p = 0.005). These results include all 40 participants. Abbreviations: ret = category retrieval; recog = recognition. Note: dotted line represents chance level of 0.5

Paired-sample, two-tailed t-tests revealed a significant drop in the frequency of sure responses from 30-minute (*M* = 27.1%, *SD* = 14.2%) to 24-hour (*M* = 21.6%, *SD* = 15.3%) retrieval (*t*(39) = 5.41, p < 0.001). In contrast, the frequency of guess responses rose from 30-minute (*M* = 39.8%, *SD* = 15.6%) to 24-hour (*M* = 50.8%, *SD* = 20.07%) category retrieval (paired t-test, two-tailed, *t*(39) = −5.36, p < 0.001). The frequency of unsure responses decreased between these retrieval time-points (paired t-test, two-tailed, *M*_30min_ = 33.1%, *SD*_30min_ = 13.4%, *M*_24h_ = 27.6% *SD*_24h_ = 15.8%, *t*(39) = 3.07, p = 0.003). On the recognition task given at 24 hours following the category retrieval, paired t-tests revealed that the frequency of sure responses almost doubled from the category retrieval at 24-hours to the recognition task (*M*_recog_ = 44.9%, *SD*_recog_ = 22.9%, *M*_24h_ = 21.6%, *SD*_24h_ = 15.3%, paired t-test, two-tailed, *t*(39) = −10.778, p < 0.001). This suggests that many of the face-object combinations that were no longer consciously accessible on the category retrieval regained conscious access with the additional object cue. Conversely, the number of guess responses decreased by half (paired t-test, two-tailed, *M*_24h_ = 50.8% *SD*_24h_ = 20.07%, *M*_recog_ = 27.1%, *SD*_recog_ = 17.3%, *t*(39) = 9.63, p < 0.001), while the number of unsure responses remained unchanged (paired t-test, two-tailed, *M*_24h_ = 27.6% *SD*_24h_ = 15.8%, *M*_recog_ = 28.0%, *SD*_recog_ = 15.2% *t*(39) = −0.15, p = 0.88). For a graphical illustration of the relative frequencies of response categories, see Figure 2A.

The retrieval accuracy on the 30-minute and 24-hour category retrieval was almost at ceiling level for the sure responses (*M*_30min_ = 87.8%, *SD*_30min_ = 10.1%; *M*_24h_ = 89.6%, *SD*_24h_ = 13.5%) but not better than chance for the guess responses (*M*_30min_ = 50.1 %, *SD*_30min_ = 10%; *μ* = 50%, one-sample, two tailed t-test, *t*(39) = 0.05, p = 0.95; *M*_24h_ = 49.4%, *SD*_24h_ = 7.1%; *μ* = 50%, one-sample, two tailed t-test, *t*(39) = −0.4, p = 0.64, see Figure 2B). The unsure responses yielded an accuracy above chance level at the 30-minute category retrieval (*M*_30min_ = 56.8%, *SD*_30min_ = 10.3%; *μ* = 50%, one-sample, two tailed t-test, *t*(39) = 4.13, p = 0.0002) and at the 24-hour category retrieval (*M*_24h_ = 54.1%, *SD*_24h_ = 9.5%; *μ* = 50%, one-sample, two tailed t-test, *t*(39) = 2.7, p = 0.01). The recognition accuracy for sure responses was at ceiling level (*M*_recog_ = 94%, *SD*_recog_ = 12.9%), significantly better than chance for the unsure responses (*M*_recog_ = 77.2%, *SD*_recog_ = 12.5%; one-sample, two tailed t-test, *μ* = 50%, *t*(39) = 14.1, p < 0.0001), and significantly better than chance for the guess responses (*M*_recog_ = 57.5%, *SD*_recog_ = 14.1%; one-sample, two tailed t-test, *μ* = 50%, *t*(39) = 3.35, p = 0.005, see Figure 2B). The Bayes Factor provided moderate evidence in favor of the alternative model for the guess responses (BF_10_ = 7.4). Hence, even when participants were only guessing which of two objects was presented along with the face during learning, their intuition guided them with a better-than-chance accuracy. These analyses included all 40 participants.

### fMRI results from univariate analyses

For the fMRI analyses, the correct guess responses were contrasted to the incorrect guess responses based on the hypothesis that a reactivation of the putative memory trace underlying a forgotten memory (therefore a guess response) would more often occur in trials that yield a correct than incorrect response. Because sure responses were almost always correct, remembered trials were also contrasted with incorrect guess responses. For brain-behavior correlations, we used the z-scored percentage correct of all guess responses (guessing accuracy) for contrasts of guess responses and the z-scored absolute number of correct sure responses (sure accuracy) for contrasts of sure responses.

### Common hippocampal areas mediated correct sure and correct guess responses at the 30-minute and 24-hour category retrieval

We asked whether forgotten memories, i.e., guess responses, were accompanied by activity increases within the episodic memory network like consciously accessible memories, i.e., sure responses (*54–58*). This was indeed the case, as revealed by the following analyses. We isolated activity increases underlying associative retrieval from activity increases underlying mere face recognition and retrieval effort by computing the contrast correct sure responses versus incorrect guess responses and the contrast correct guess responses versus incorrect guess responses. These contrasts were then statistically conjugated to find common brain activation underlying correct sure and correct guess responses.

For the 30-minute category retrieval, the conjunction analysis using the data from the small FOV fMRI sequence yielded results in the right hippocampus (right hippocampus: p_uncor_ < 0.001, *T(285)* = 4.05, peak at coordinates based on the Montreal Neurological Institute (MNI) [30,−32,−9], see Figure 3A and Table S2). Further significant conjunctive results were located in the left posterior cingulate gyrus. These results were not replicated in the whole-brain fMRI sequence that had a slightly worse spatial resolution: the same conjunction analysis with data from the whole-brain fMRI sequence yielded no significant result at p < 0.001 uncorrected and k = 10 voxel extent.

**Figure 3.**
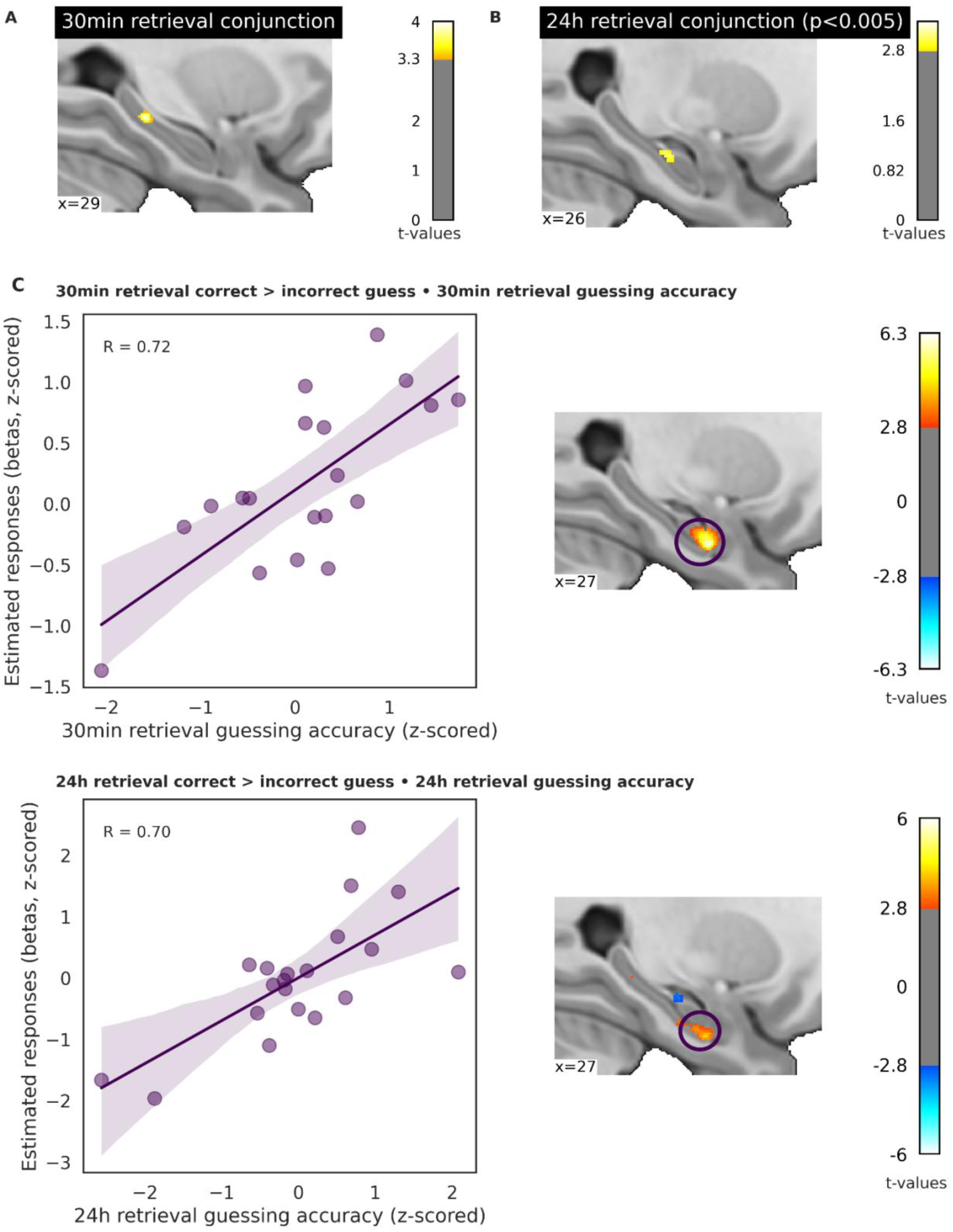
Right hippocampal activity increased for both correct sure and correct guess responses at the 30-minute and the 24-hour category retrieval and correlated for guess responses with guessing accuracy at both testing time points. A) Common right hippocampal activity increases underlying correct sure and correct guess responses at the 30-minute category retrieval. We isolated activity increases underlying associative retrieval from activity increases underlying mere face recognition, task characteristics, and retrieval effort by computing the contrast correct sure responses > incorrect guess responses and the contrast correct guess responses > incorrect guess responses. These contrasts were then statistically conjugated to find common brain activation underlying correct sure and correct guess responses. For a table of cluster sizes and t-values, see Table S2. Results presented in this panel stem from the small FOV fMRI sequence. B) Common right hippocampal activity rises underlying correct sure and correct guess responses at the 24-hour category retrieval. Statistical analysis analogous to 3A. For a table of cluster sizes and t-values, see Table S3. Results presented in this panel stem from the small FOV fMRI sequence. C) Right hippocampal activity underlying correct guess responses on the category retrieval correlated with the guessing accuracy on the category retrieval at 30 minutes and at 24 hours. Upper panel: positive brain-behavior correlation in the right hippocampus. We show the contrast of correct guess > incorrect guess responses correlated with the guessing accuracy on the category retrieval at 30 minutes (for cluster sizes and t-values, see Table S4). Lower panel: positive brain-behavior correlation in the right hippocampus. We show the contrast of correct guess > incorrect guess responses correlated with the guessing accuracy on the category retrieval at 24-hours (for cluster sizes and t-values, see Table S5). Outliers (above absolute z-value of 3) were removed. Results presented in this panel stem from the small FOV fMRI sequence.

For the 24-hour category retrieval, the conjunction analysis using the data from the small FOV fMRI sequence yielded also a result in the right hippocampus, albeit slightly below our chosen threshold of p < 0.001 uncorrected (right hippocampus: p_uncor_ = 0.0012, *T(285)* = 3.07, peak at MNI [26,−18,−17]). For illustrative purposes, see Figure 3B at p < 0.005 uncorrected. Significant results (at p < 0.001) were located in the left lingual gyrus and bilateral posterior cingulate gyrus (see Table S3). The same conjunction analysis computed with data from the whole-brain fMRI sequence yielded no significant result. The lower power of conjunctive activity in the right hippocampus head at 24 hours might be due to a distinct overnight consolidation for consciously accessible versus consciously inaccessible memories, see below.

Hence, the hippocampus and other relevant structures of the episodic memory retrieval network were involved in the reactivation of both consciously accessible and consciously inaccessible associative memories at the 30-minute and at the 24-hour category retrieval.

### Hippocampal activity underlying correct guess responses correlated with the guessing accuracy on the category retrievalat 30 minutes and at 24 hours

Given that correct guess responses and correct sure responses were accompanied by activations in the same right hippocampal region, we then examined whether right hippocampal activity underlying correct guess responses versus incorrect guess responses would correlate with guessing accuracy across participants. Using the small FOV fMRI sequence for this analysis, a cluster in the right hippocampus head correlated with the guessing accuracy and even survived family-wise error correction (peak at MNI [27,−8,−25], cluster-level p_FWE_ = 0.019, T(18) = 6.31, R = 0.72, see Figure 3C, upper panel and Table S4 for all correlation results). Besides the right hippocampus, significant correlations were located in multiple bilateral inferior and middle temporal regions putatively supporting the reactivation of semantic object-related information and the precuneus and the posterior cingulate gyrus related to visual imagery.

The same correlation computed for the 24-hour category retrieval using the small FOV fMRI sequence yielded a significant cluster in the same area within the right hippocampus head (peak at MNI [28, −10,−28], p_uncor_ > 0.001, T(18) = 4.70, R = 0.70, see Figure 3C, lower panel) besides multiple further clusters located in the right and left hippocampus (see Table S5). This result suggests that the bilateral hippocampus is involved in the retrieval of consciously inaccessible memories at 24 hours following learning, probably due to over-night memory consolidation, as detailed in a section below (see also Fig. 5). Besides bilateral hippocampal areas, significant correlations were located in several bilateral middle temporal regions probably supporting the reactivation of semantic object-related information.

### Hippocampal functional connectivity during learning and 30-minute category retrieval correlated with the 30-minute guessing accuracy for forgotten associations

We next asked which neural circuits would interact with the right hippocampus when forming new associative memories that would later be forgotten. To this end, we computed a general psycho-physiological interaction analysis (gPPI) with the right hippocampus head as the seed region (based on the correlative results above) using the data of the whole-brain fMRI sequence. The comparison of learning trials that would yield correct versus incorrect guess responses at the 30-minute category retrieval (subsequent memory analysis) revealed that seed-based connectivity estimates correlated between-subjects with guessing accuracy. This was true for the right hippocampus head-to-bilateral anterior cingulate/medial superior frontal gyrus connectivity (peak at MNI [12,42,−4], peak-level p_FWE_ < 0.05, *T(18)* = 7.8, R = 0.92, see Figure 4A) and right hippocampus head-to-left hippocampus, right hippocampus head-to-bilateral lingual gyrus, and right hippocampus head-to-bilateral cuneus/precuneus connectivity (all results listed in Table S6). When computing this same correlation analysis with data collected during the 30-minute category retrieval, the right hippocampus-to-medial superior frontal gyrus connectivity again correlated with guessing accuracy (peak at MNI [12,46,28], cluster-level p_FWE_ < 0.001, *T(18)* = 6.25, R = 0.85, see Figure 4B). Connectivity from right hippocampus to bilateral lingual gyrus, right cuneus and bilateral precuneus no longer correlated at the 30-minute category retrieval. However, right hippocampus-to-various prefrontal regions and right hippocampus-to-left inferior temporal gyrus connectivity correlated between-subjects with retrieval accuracy at the 30-minute category retrieval (all results listed in Table S7). No significant correlations were found between seed-based connectivity and guessing accuracy at 24-hour category retrieval. The reason for this lack of significant results at 24 hours might be the overnight consolidation of forgotten memories that provided for a deeper rooting of forgotten memories in the hippocampus rather than the neocortex (this will be detailed below).

**Figure 4:**
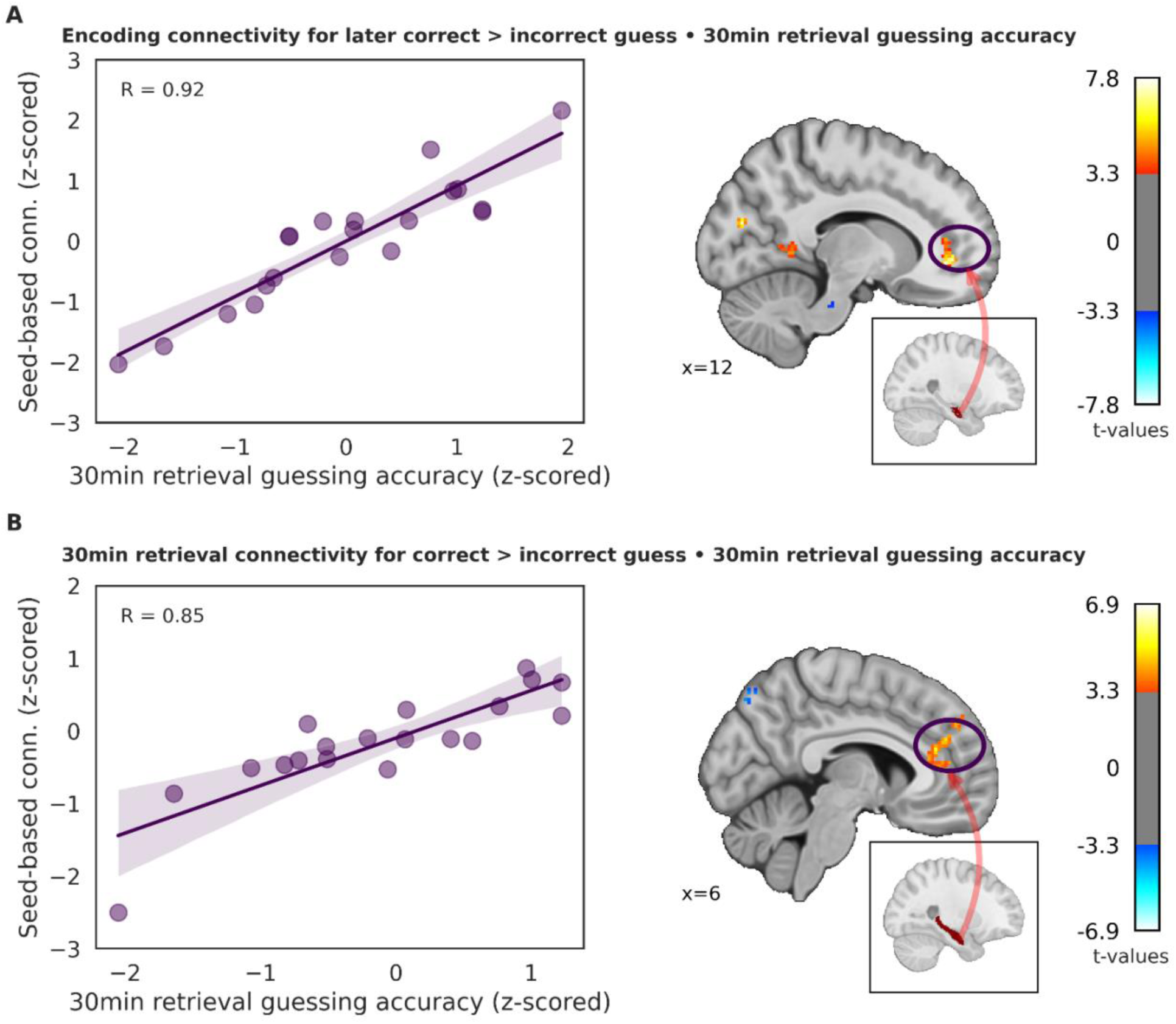
Hippocampal functional connectivity measured during learning and during the 30-minute category retrieval correlated with the 30-minute category retrieval guessing accuracy. A) Hippocampal-medial prefrontal functional connectivity during learning correlated with the later recorded 30-minute category retrieval guessing accuracy. We computed the functional connectivity between the seed region in the right hippocampus head (see box inlet) with neocortical regions during learning using a gPPI. In participants with a higher guessing accuracy on the 30-minute category retrieval, the functional connectivity was stronger between the right hippocampus head and the right anterior cingulate gyrus and the right medial prefrontal cortex. Outliers (above absolute z-value of 3) were removed. For a table of cluster sizes and t values see Table S6. Results presented in this panel stem from the whole-brain fMRI sequence. B) Hippocampal-medial prefrontal functional connectivity during the 30-minute category retrieval correlated with the 30-minute category retrieval guessing accuracy. We computed the functional connectivity between the seed region in the right whole hippocampus (see box inlet) with neocortical regions during the 30-minute category retrieval using a gPPI. Participants with a higher guessing accuracy on the 30-minute category retrieval, exhibited a stronger functional connectivity between the right hippocampus and the right medial prefrontal cortex. Outliers (above absolute z-value of 3) removed. For cluster sizes and t-values see Table S7. Results presented in this panel stem from the whole-brain fMRI sequence.

### Overnight consolidation brought a neocorticalization for sure responses and a deeper/broader hippocampal implementation for guess responses

Following the first visit to the MR Center, the learned face-object associations underwent overnight consolidation before participants reactivated the associations again during the category retrieval on day 2. Following overnight consolidation, memory traces underlying sure versus guess responses became implemented more strongly in the neocortex as the standard model of memory consolidation predicts. This was the result of the stacked contrast (24-hour category retrieval: correct sure > correct guess) > (30-minute category retrieval: correct sure > correct guess responses) computed with the whole-brain fMRI data. It revealed strong overnight increases in activity in many bilateral fronto-parietal, temporal and occipital regions (all results in Table S8). The same computation with data from the small FOV fMRI sequence did not yield significant results.

In contrast, memory traces underlying correct guess responses on the category retrieval were implemented more broadly and more strongly within bilateral hippocampus on the second versus the first day. This was the result of the comparison of correct guess responses given at the 24-hour category retrieval versus correct guess responses given at the 30-minute category retrieval using the data of the small FOV fMRI sequence. The result revealed overnight activity increases in bilateral hippocampus head (left hippocampus: peak at MNI [−27, −16, −14], p_uncor_ < 0.001, *T(18)* = 4.62, right hippocampus: peak at MNI [39,−22,−12], p_uncor_ < 0.001, *T(18)* = 3.91, see Table S9 for all results). The reverse contrast did not yield significant results in the hippocampus but results in the left-hemisphere: entorhinal area and middle temporal gyrus (see Table S10). Repeating this analysis using the data from the whole-brain fMRI sequence yielded merely one cluster in the inferior occipital gyrus for correct guess responses at 24 hours versus correct guess responses at 30 minutes (peak at MNI [−30 −90 −12], p_uncor_ < 0.001, *T(18)* = 3.81).

Next, we computed a gPPI analysis with the seed region in the right hippocampus head using the small FOV fMRI sequence to examine overnight changes in the functional connectivity within the medial temporal lobe underlying correct guess responses. Contrasting the functional connectivity underlying the correct guess responses given at the 24-hour category retrieval with the functional connectivity underlying correct guess responses given at the 30-minute category retrieval and correlating the results with the 24-hours guessing accuracy, we see an overnight increase in the behaviorally relevant functional connectivity of the right hippocampus head with left hippocampal regions (peak at MNI [−26,−30,−4], cluster-level p_FWE_ = 0.011, *T(18)* = 5.37, R = 0.61), with right entorhinal cortex, and with several regions within the right lateral temporal lobe (Figure 5; for all results, see Table S11). Hence, the overnight consolidation left the memory traces underlying guess responses implemented more broadly and deeply in bilateral hippocampus, which benefitted the accuracy of the guess responses given at 24 hours.

**Figure 5.**
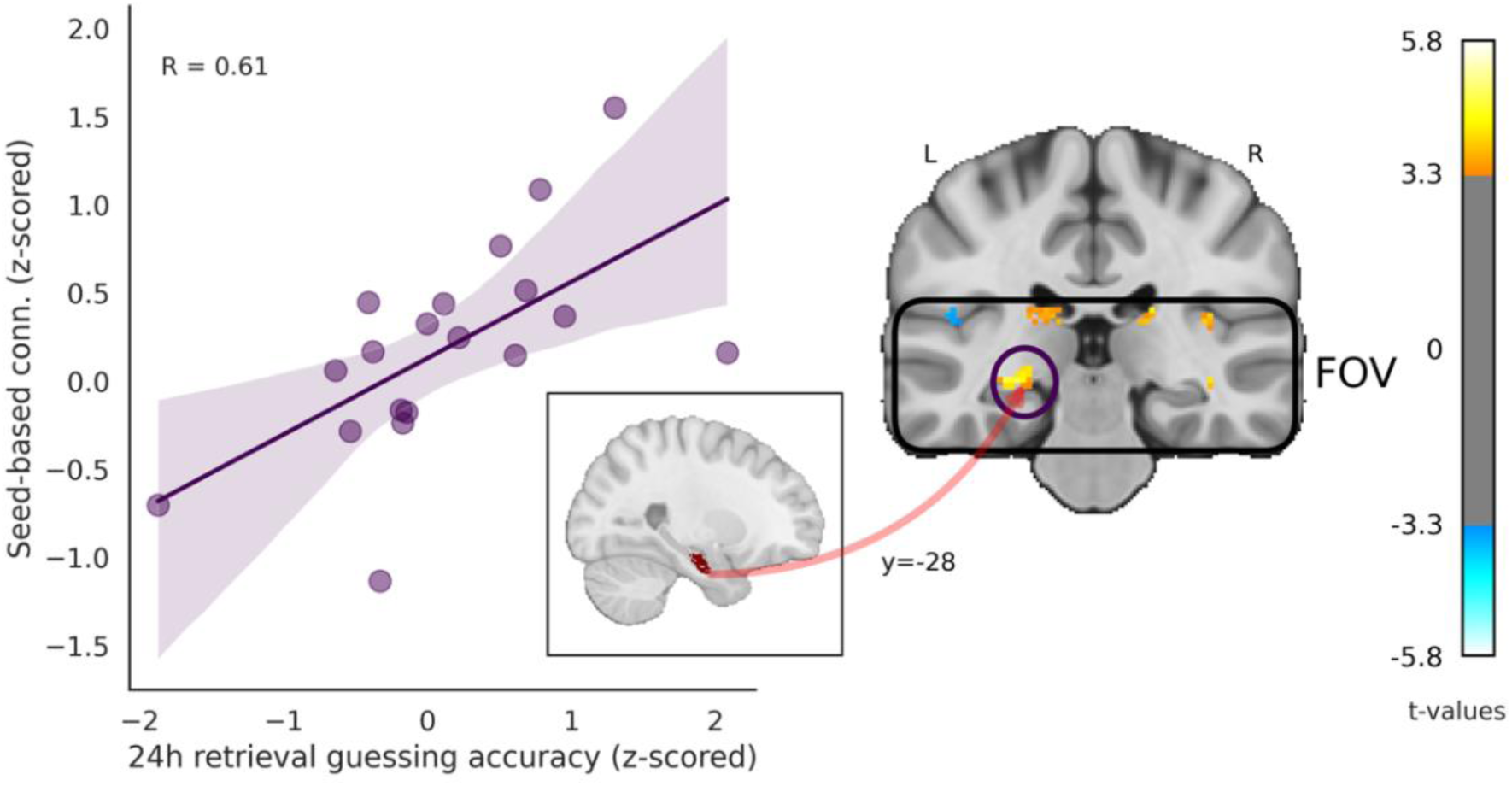
Overnight right-left hippocampal functional connectivity increases improved guessing accuracy at the 24-hour category retrieval. We computed a gPPI analysis with the seed region in the right hippocampus head (see box inlet) using the small FOV fMRI sequence to examine overnight changes in the functional connectivity within the medial temporal lobe underlying correct guess responses on the category retrieval. We contrasted the functional connectivity underlying the correct guess responses given on the 24-hour category retrieval with the functional connectivity underlying correct guess responses given on the 30-minute category retrieval and then correlated the result with the 24-hour guessing accuracy on the category retrieval. Connectivity estimates and guessing accuracy were z-scored. Outliers (above absolute z-value of 3) removed. See Table S11 for all results.

### Medial prefrontal cortex activation during guess responses given at the 24-hour category retrieval predicted the subsequent regain of conscious access to the memories

As mentioned in the behavioral results section, the mean guessing accuracy on the 24-hour category retrieval was at chance level but then rose significantly above chance level on the immediately following recognition task, where the face plus two objects (original object and foil object) were presented along with the face cue as additional retrieval cues. Hence, the addition of a valid object retrieval cue to the face retrieval cue (which was also present for category retrieval) improved the guessing accuracy and sometimes even allowed participants to regain conscious access to the previously inaccessible memories (see Figure 2A). Because the recognition task immediately followed the 24-hour category retrieval, we were wondering which brain regions elevated activation levels during the 24-hour category retrieval and whether these activation elevations would predict the subsequent regain of conscious access during the recognition. To this end, we contrasted the guess responses on the 24-hour category retrieval that would subsequently yield correct sure responses (i.e., conscious access) on the recognition task with those other guess responses given on the 24-hour category retrieval that would yield correct guess responses (i.e., no conscious access) on the recognition. We used the data from the whole-brain fMRI sequence. The medial prefrontal cortex was strongly activated for later correct sure versus correct guess responses given on the recognition task (peak at MNI [0 30 −16], p_uncor_ < 0.001, *T(18)* = 4.19, see Figure S1A and Table S12). This suggests that consciously inaccessible memories (guesses) on the 24-hour category retrieval that possess (versus not) a medial prefrontal engram component, tend to become consciously accessible again when given an additional retrieval cue (the object).

### Additional object cues given on the recognition task permitted the reactivation of engram components in the episodic memory network underlying sure responses and guess responses

Because the right hippocampus head was involved in both guess and sure responses during the category retrieval, we asked which neural circuits the right hippocampus head would engage in for successful face-object recognition. We computed a gPPI analysis with the seed region in the right hippocampus head using the whole-brain fMRI sequence to examine the functional connectivity underlying guess and sure responses given during recognition. Like in earlier analyses, we subtracted the functional connectivity for incorrect guess responses from the functional connectivity for correct sure and for correct guess responses given during recognition. The results were correlated with the sure and guess recognition accuracy, respectively. Then, we computed a conjunction analysis on the two gPPI results to determine commonly activated and behaviorally relevant neural circuits underlying recognition. The functional connectivity of the right hippocampus head with bilateral lingual gyrus and bilateral calcarine cortex was crucial for recognition success (peak at MNI [−4 −82 −2], p_uncor_ < 0.001, *T(18)* = 3.53, note that this was the only cluster found for this conjunction analysis). Hence, the functional connectivity of the right hippocampus with areas of the visual cortex was relevant for the explicit (remembered associations) and implicit (forgotten associations) recognition of the previously learned face-object combinations.

Zooming in on the forgotten associations, we were wondering about the difference in the behaviorally relevant functional connectivity of the right hippocampus head for those forgotten memories that still yielded correct versus incorrect guess responses on the recognition. We computed a gPPI analysis with data from the whole-brain fMRI sequence for the seed region in the right hippocampus head comparing correct versus incorrect guess responses on the recognition task and correlated the results between-subjects with the individual accuracy of the guess responses. The accuracy of the guessed recognition responses increased with a connectivity increase between the right hippocampus head and the left hippocampus tail, between the right hippocampus head and bilateral lingual gyrus, between the right hippocampus head and the left middle temporal gyrus as well as between the right hippocampus head and a large bilateral fronto-parietal network (all results in Table S13).

Unlike face-cued relational retrieval during the category retrieval, relational (face-object) recognition may in principle be mediated by at least three distinct memory mechanisms: episodic memory, priming, and familiarity. While the feeling of familiarity is consciously accessible and therefore cannot have influenced guess responses, priming is unconscious and might have added to the correct guess responses given on the recognition task (*59*, *60*). Hence, perceptual and/or semantic associative priming may have contributed to the above chance guessing result on the recognition. Nevertheless, guessing accuracy on the recognition task was mediated by bilateral hippocampal connectivity, which suggests that guessing accuracy was also driven by episodic memory.

Zooming in on the consciously accessible associations, we computed the same gPPI analysis for correct sure versus incorrect guess responses given on the recognition task and correlated the result between-subjects with the individual accuracy of the sure responses. The result was a similar connectivity of the right hippocampus head with bilateral lingual gyrus, with the right middle temporal gyrus as well as a large bilateral fronto-parietal network, but no connectivity with the left hippocampus tail (see Table S14 for all results).

Hence, response accuracy during guessing was mediated by bilateral hippocampal connectivity. Response accuracy during both guessing and sure responses was mediated by the connectivity of the right hippocampus head with visual cortices and a large bilateral fronto-parietal network. It thus appears that the addition of the relevant object cue during recognition allowed for a better reactivation of engram components in the episodic memory network underlying both sure and guess responses. Note that the right hippocampus head-to-left hippocampus tail connectivity result for guess responses was not replicated when using the small FOV fMRI sequence. The cause might be participant differences because of the between-subjects design in implicit retrieval or a poor signal in this very posterior left hippocampal region.

### fMRI results from representational similarity analyses

We asked whether more fine-grained engram components, namely individual voxel patterns, would reactivate during distinct memory stages and reflect the persistence or shift regarding certain engram components over time. We expected the repetition of voxel patterns underlying memories in those brain areas that yielded significant results in the above reported univariate analyses. Therefore, we chose a region-of-interest approach deriving the regions-of-interest from the above reported univariate results and from the priori hypothesis that the hippocampal engram components would persist when a memory becomes consciously inaccessible. Of particular interest regarding the trajectory of episodic memories were recurring or shifting voxel patterns in the hippocampus. We computed representational similarity analyses (RSA), i.e., pair-wise comparisons of voxel patterns, comparing encoding – retrieval similarity (ERS) for the 30-minute and 24-hour category retrieval, encoding – recognition (at 24 hours) similarity, and retrieval (30 minutes) – retrieval (24 hours) similarity (RRS) for the category retrieval. We separated trials into guess and sure responses to compare the trajectory of engram components for consciously accessible and consciously inaccessible memories. By subtracting RSA values underlying incorrect guess responses from RSA values underlying correct guess and correct sure responses, we isolated estimates due to associative retrieval success by eliminating estimates that originate from the mere repetition of face cues (yielding face recognition) and by eliminating estimates that originate from shared task requirements (STAR Methods). Thus, the below reported similarity estimates underly successful associative encoding and associative retrieval operations. The height threshold in all RSA analyses was set to p < 0.05, uncorrected for multiple comparisons. We report a Bayes factor for each t-test and Pearson correlation.

### Similarity of voxel patterns between encoding and the 30-minute category retrieval were relevant for retrieval accuracy

Using data from the small FOV fMRI sequence, voxel pattern similarity in the left hippocampal body between encoding and the 30-minute category retrieval for correct sure responses tended towards a significant correlation with the sure accuracy across subjects (p = 0.086, R = 0.39, BF_10_ = 1.1). The same computation yielded neither a tendency nor a significant result in the medial temporal lobe for correct guess responses.

Within neocortex (using data from the whole-brain fMRI sequence), voxel patterns were significantly similar in the bilateral inferior temporal gyrus for correct sure responses (*t(16*) = 2.3, p = 0.035, BF_10_ = 1.9, see Figure S2), with no significant between-subjects correlation of the similarity values with the sure accuracy. For correct guess responses, this computation revealed non-significant mean value comparisons but a significant between-subjects correlation between the degree of similarity of voxel patterns in the bilateral middle temporal gyrus and guessing accuracy (p = 0.022, R = 0.55, BF_10_ = 3.3, see Figure 6A). It thus appears that the reactivation of engram components in the inferior and middle temporal gyrus contributed to the conscious and unconscious reactivation of memories at 30 minutes following encoding.

**Figure 6.**
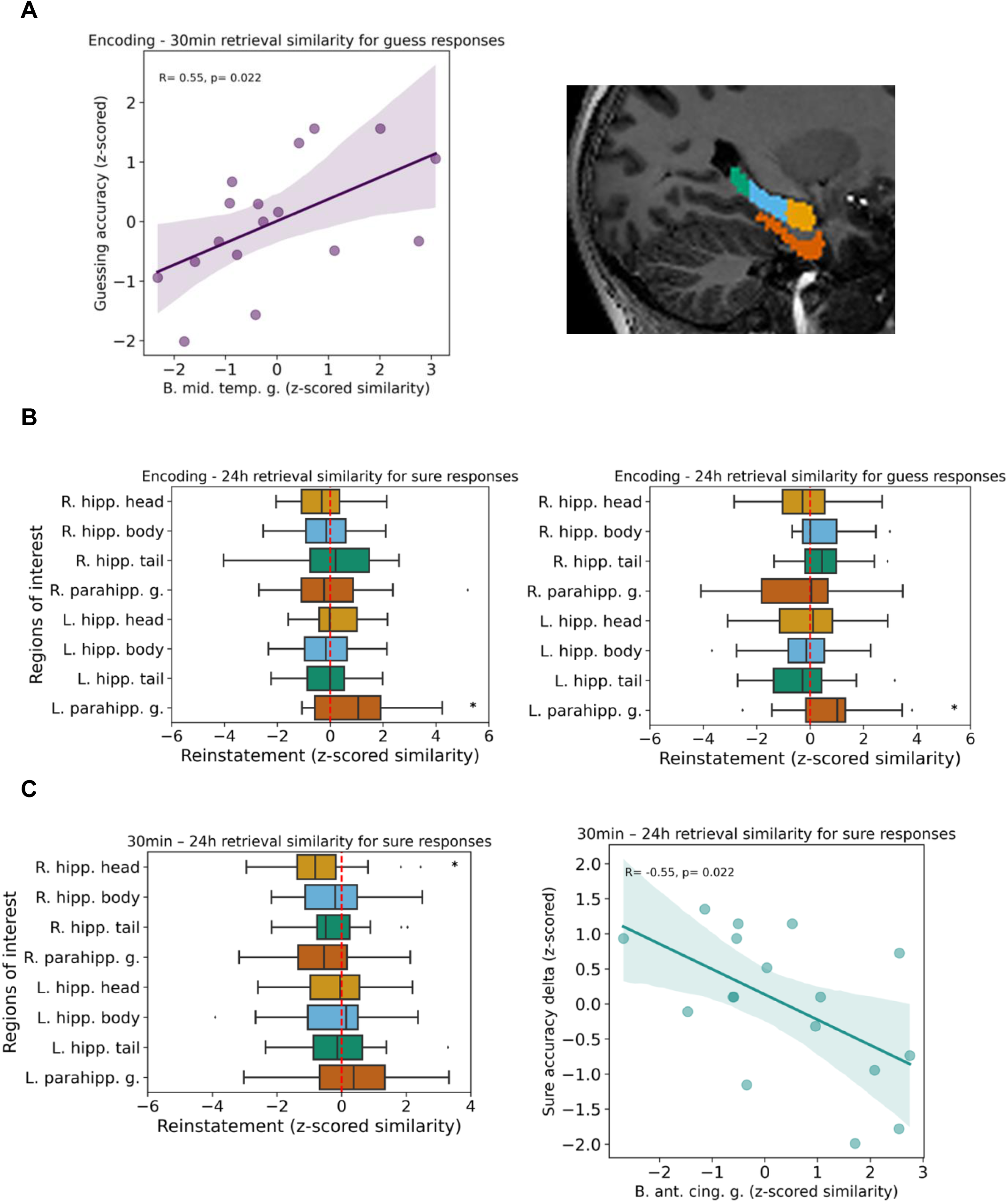
Encoding – retrieval and retrieval – retrieval similarity. A) Left panel: Similarity of voxel patterns between encoding and the 30-minute category retrieval were relevant for retrieval success. Significant between-subjects correlation of encoding-retrieval similarity values with guessing accuracy on the 30-minute category retrieval in bilateral middle temporal gyrus. Results presented in this panel were acquired with the whole-brain fMRI sequence. Right panel: small FOV ROI overview for B) and C) B) Similarity of voxel patterns between encoding and the 24-hour category retrieval were relevant for retrieval success. Encoding-retrieval similarity values were significant in left parahippocampal gyrus for both correct guess responses (left) and correct sure responses (right) (t(19) = 2.65, p = 0.015, BF_10_ = 3.5; t(19) = 2.35, p = 0.029, BF_10_ = 2.1, respectively). Results presented in this panel were acquired with the small FOV fMRI sequence. C) Dissimilarity of voxel patterns between the 30-minute and the 24-hour category retrieval were relevant for retrieval success. Left panel: 30-minute – 24-hours category retrieval dissimilarity in the right hippocampus head for correct sure responses. Results presented in this panel were acquired with the small FOV fMRI sequence. Right panel: 30-minute – 24-hours category retrieval dissimilarity values correlated between-subjects with the delta of correct sure responses (24-hours sure accuracy minus 30-minute sure accuracy) in bilateral anterior cingulate gyrus (B. ant. cing. g.). Results presented in this panel were acquired with the whole-brain fMRI sequence. *p < 0.05, **p < 0.01 by Student’s t test. Abbreviations: ROI, region of interest; R., right; L., left; hipp., hippocampus; parahipp., parahippocampus; g., gyrus.

### Similarity of voxel patterns between encoding and the 24-hour category retrieval were relevant for retrieval success

Following the first visit at the MR Center, the encoded face-object associations underwent overnight consolidation before participants reactivated the associations again during their second visit at the MR Center on day 2. Overnight memory consolidation may have stabilized the activated voxel patterns from their inception to their 24-hour reactivation during the category retrieval. For both correct guess (p = 0.029, *t(19)* = 2.35, BF_10_ = 2.1) and correct sure (p = 0.015, *t(19)* = 2.65, BF_10_ = 3.5) responses, voxel patterns were significantly similar between encoding and the 24-hour category retrieval within the left parahippocampal gyrus (see Figure 6B). Moreover, the degree of this left parahippocampal pattern similarity for sure responses tended to correlate between-subjects with the sure accuracy on the 24-hour category retrieval (p = 0.058, R = 0.43, BF_10_ = 1.5); the same was true for the left hippocampal body (p = 0.059, R = 0.44, BF_10_ = 1.5). Note that the pattern similarity in the left hippocampal body had already correlated between subjects with the sure accuracy for the encoding – 30-minute comparison. Hence, medial temporal pattern reinstatements were behaviorally relevant for the conscious retrieval at both time-points. For correct guess responses, however, correlations between medial temporal encoding-retrieval pattern reinstatements and retrieval success did not reach significance. A potential reason could be consolidation: the time after encoding appeared to have strengthened and transformed the medial temporal engram components of forgotten memories. Results from this paragraph are based on data from the small FOV fMRI sequence.

Within neocortex, the similarity of voxel patterns between encoding and the 24-hour category retrieval underlying correct sure responses revealed non-significant mean value comparisons but a significant between-subjects correlation of the similarity values with the sure accuracy within the bilateral middle temporal gyrus (p = 0.006, R = 0.64, BF_10_ = 10.4, see Figure S3) as previously seen for the correctly guessed trials for the encoding – 30-minute category retrieval comparison. The similarity of voxel patterns between encoding and 24-hour category retrieval also revealed non-significant mean value comparisons for correct guess responses but a significant between-subjects correlation with guessing accuracy in bilateral cuneus (p = 0.049, R = 0.48, BF_10_ = 1.8) plus non-significant correlations within bilateral inferior temporal sulcus (p = 0.099, R = −0.41, BF_10_ = 1.0) and bilateral anterior cingulate gyrus (p = 0.100, R = −0.41, BF_10_ = 1.0).

### (Dis)similarity of voxel patterns between the 30-minute and the 24-hour category retrieval were relevant for retrieval success

The retrieval-retrieval comparison of voxel patterns, i.e., the comparison between the 30-minute and the 24-hour category retrieval, for correct sure responses yielded a significant dissimilarity for the right hippocampus head (p = 0.035, *t(19)* = −2.26, BF_10_ = 1.8, see Figure 6C, left panel). The same analysis for correct guess responses revealed also a significant dissimilarity in the right hippocampus, however in its tail region (p = 0.046, *t(19)* = −2.12, BF_10_ = 1.5, not displayed). On the other hand, there was a retrieval-retrieval similarity (not dissimilarity) of voxel patterns in the right hippocampus head underlying correct guess responses that correlated significantly (p = 0.018, R = 0.52, BF_10_ = 3.7, see Figure S4) between-subjects with the delta of correct guess responses (i.e., guessing accuracy at 24 hours minus the guessing accuracy at 30 minutes). Hence, participants exhibiting a similar right hippocampal retrieval-retrieval voxel pattern yielded a better retrieval accuracy at 24 hours versus 30 minutes. Although a right hippocampal pattern dissimilarity hallmarked correct retrieval responses at 24 hours versus 30 minutes for both sure and guess responses, a pattern similarity within the right hippocampus head still aided a few participants, who lacked conscious access to memories at both retrieval time-points. Results from this paragraph are based on data from the small FOV fMRI sequence.

Within neocortex, the (dis)similarity of voxel patterns between the 30-minute and the 24-hour category retrieval underlying correct sure responses revealed non-significant mean value comparisons but a significant between-subjects correlation of pattern dissimilarity with the delta of correct sure responses within bilateral anterior cingulate gyrus (p = 0.021, R = −0.55, BF_10_ = 3.4, see Figure 6C, right panel). Thus, the overnight memory consolidation may have also changed the anterior cingulate engram components – and not just the right hippocampal engram components – such that a pattern dissimilarity supported a successful retrieval on the next day. For correct guess responses, the (dis)similarity of voxel pattern between the 30-minute and the 24-hour category retrieval revealed also non-significant mean value comparisons, yet a significantly positive between-subjects correlation of pattern similarity with the delta of guessing accuracy within bilateral cuneus (p = 0.005, R = 0.64, BF_10_ = 10.8, not displayed). Hence, participants with a similar cuneal retrieval-retrieval voxel pattern yielded a better retrieval accuracy at 24 hours versus 30 minutes. Therefore, also the cuneal pattern similarity appears to have aided a few participants, who lacked conscious access to memories at both retrieval time-points. Overall, these results show that, especially for dynamic engram components such as the hippocampus and anterior cingulate gyrus, behaviorally relevant dissimilarity can occur.

### Encoding – recognition pattern similarity

Unlike the retrieval-retrieval dissimilarity results presented above, there was not a single pattern dissimilarity result for the encoding-recognition comparison. For correct sure responses on the recognition task, voxel patterns were significantly similar between encoding and recognition within the right hippocampal body (p = 0.001, *t(19)* = 3.9, BF_10_ = 34.0) and the right parahippocampal gyrus (p = 0.048, *t(19)* = 2.12, BF_10_ = 1.5, Figure 7A), with no significant between-subjects correlations of pattern similarity with recognition accuracy. For correct guess responses on the recognition task, neither mean value comparisons nor correlations between pattern reinstatements and recognition accuracy reached significance. Results from this paragraph are based on data from the small FOV fMRI sequence.

**Figure 7.**
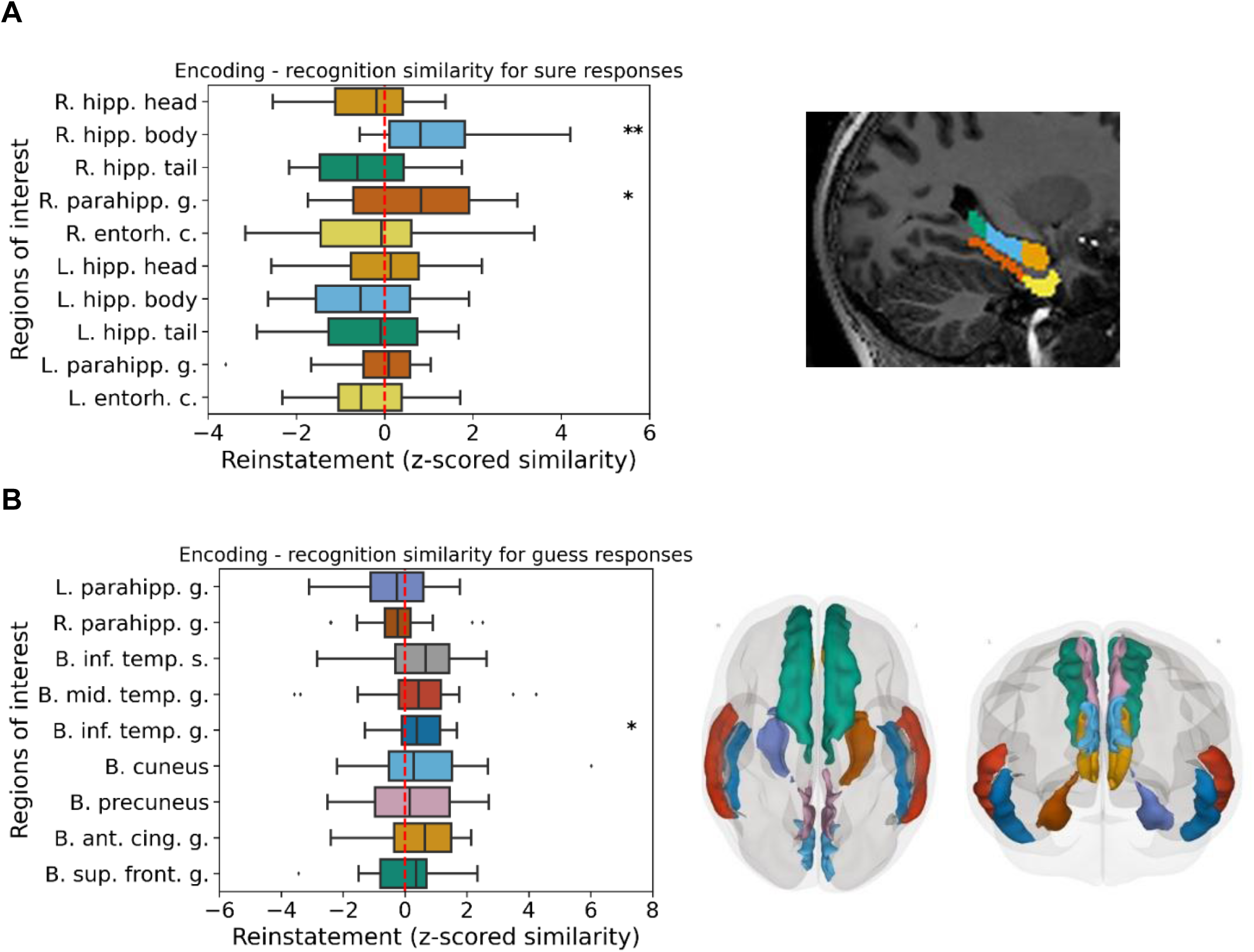
Encoding – recognition pattern similarity. A) Encoding-recognition pattern similarity in the right hippocampal body and right parahippocampal gyrus for correct sure responses on the recognition task (t(19) = 3.91, p = 0.001, BF_10_ = 34.0; t(19) = 2.11, p = 0.048, BF_10_ = 1.5, respectively). Results presented in this panel were acquired with the small FOV fMRI sequence. B) Encoding-recognition pattern similarity in bilateral inferior temporal gyrus for correct guess responses on the recognition task (p = 0.038, t(17) = 2.25, BF_10_ = 1.8). Results presented in this panel were acquired with the whole-brain fMRI sequence. Right panels: ROI overview. *p < 0.05, **p < 0.01 by Student’s t test. Abbreviations: R., right; L., left; B., bilateral; parahipp., parahippocampus; hipp., hippocampus; entorh., entorhinal; inf., inferior; mid., middle; ant., anterior; cing., cingulate; sup., superior; front., frontal; s., sulcus, g., gyrus c., cortex.

Within neocortex, the similarity of voxel patterns between encoding and recognition underlying correct sure responses on the recognition task revealed a significant mean value comparison in the bilateral inferior temporal sulcus (p = 0.020, *t(17)* = 2.55, BF_10_ = 2.9, see Figure S5). The degree of this inferior temporal pattern similarity did not correlate with recognition success. Moreover, a pattern similarity in bilateral anterior cingulate gyrus correlated significantly between-subjects with recognition accuracy (p = 0.042, R = 0.48, BF_10_ = 2.0, see Figure S6) but was insignificant in terms of mean value comparison. Also, the similarity of voxel patterns between encoding and recognition for correct guess responses revealed a significant mean value comparison in the bilateral inferior temporal gyrus (p = 0.038, *t(17)* = 2.25, BF_10_ = 1.8, see Figure 7B). The degree of this inferior temporal pattern similarity did not correlate with accuracy of guess responses given on the recognition. Overall, these results show that correct sure responses were accompanied by encoding – recognition voxel pattern similarity in the medial temporal lobe, consistent with an episodic reinstatement, while for guess trials, the encoding-recognition reinstatement of neocortical voxel patterns seems to be relevant for retrieval. A potential reason for the lack of encoding-recognition medial temporal similarity results for correct guess responses could be consolidation: the time after encoding appeared to have strengthened and transformed the medial temporal engram components of forgotten memories.

## DISCUSSION

Inspired by recent engram research in mice suggesting that the engrams of forgotten memories remain implemented in the brain, we asked whether engrams underlying human episodic memories would also persist and whether they influence human behavior implicitly? We mapped the fate of 96 new engrams at the systems level using high-resolution fMRI at 7 Tesla during learning, an immediate, a 30-minute, and a 24-hour category retrieval. Participants indicated on each retrieval trial, whether they remembered or forgot the memory. Almost all 96 associative memories could be formed during the one-trial learning event, and they were consciously retrieved at the immediately given category retrieval. Hence, encoding was successful. Univariate and multivariate analyses of the fMRI data revealed that the engrams of subsequently forgotten memories remained implemented in the episodic memory network and continued to influence the accuracy of guess responses at later stages of category retrieval. The engrams of forgotten memories became deeper implemented within bilateral hippocampus overnight, while consciously accessible memories were neocorticalized overnight. For consciously accessible and inaccessible (forgotten) memories, engram components within right hippocampus and anterior cingulate cortex shifted from the 30-minute to the 24-hour category retrieval. Hence, forgotten episodic memories appear to remain implemented in the episodic memory network and continue to influence human behavior implicitly.

### Traces of consciously inaccessible memories remained in the hippocampus and continued to guide retrieval choices

Although the 30-minute and the 24-hour guessing accuracies on the category retrieval were not above chance level on average, brain activity in the right hippocampus head – an essential brain region for episodic memory (*35*, *39*) – underlying correct versus incorrect guess responses correlated with guessing accuracy across participants at both time-points. This strongly suggests that the episodic memory traces persisted in the brain despite loss of conscious access to the memories and that the traces continued to influence the participants’ choice behavior at test. Memory traces underlying consciously accessible (sure responses) and consciously inaccessible (guess responses) memories overlapped in the episodic memory retrieval network. The engram-related activity was generally a bit weaker and more circumscribed for forgotten versus remembered memories, although these differences were not significant statistically. The rather weak activity underlying forgotten memories nevertheless influenced the participants’ choice behavior at retrieval (*47*, *48*, *61*).

### Distinct consolidation trajectories for consciously accessible and inaccessible memories

Interestingly, consciously accessible and consciously inaccessible memories underwent distinct consolidation trajectories overnight. Consciously accessible memories relied on neocortical structures immediately following encoding and became even more broadly neocorticalized following overnight consolidation, which is typical for systems consolidation of episodic memories. Goto and colleagues (*20*) had previously demonstrated that a first wave of long-term potentiation acts locally in the hippocampus during the minutes following learning, a second wave of hippocampal long-term potentiation-dependent consolidation occurs during the first sleep phase following learning; and a third wave of long-term potentiation occurs in the anterior cingulate gyrus on the second day following learning and supports the strengthening of neocortical engram components. It appears that the strong, consciously accessible memories soon underwent the third wave of memory consolidation because they became quickly represented in anterior cingulate gyrus and medial prefrontal cortex. At the behavioral level, the consciously accessible memories remained relatively stable overnight: virtually all consciously accessible memories at the 24-hour category retrieval were already consciously accessible at the 30-minute category retrieval.

The consciously inaccessible (forgotten) memories recruited fewer neocortical areas immediately following encoding and following overnight consolidation, rather they became more broadly implemented in bilateral hippocampus: the left and right hippocampus were more strongly activated during guess responses given at the 24-hour relative to the 30-minute category retrieval. Moreover, the inter-hippocampal connectivity became stronger overnight and correlated with the 24-hour category guessing accuracy. Therefore, overnight memory consolidation may have compensated for a weaker representational status of forgotten memories with the purpose of rescuing these memories from degradation by implementing them deeper in the medial temporal lobe and only later into the neocortex. This compensation appears to have occurred during the first two waves of memory consolidation according to Goto (*20*) and colleagues, which act within the hippocampus. Hence, the consolidation of forgotten memories proceeded slower than described by Goto et al. and was much slower than the consolidation of the consciously accessible memories in this study. Interestingly though, the stronger memories among the forgotten memories also activated the anterior cingulate gyrus at the 24-hour category retrieval, like the consciously accessible memories did, and this subgroup of forgotten memories regained conscious access during the recognition. The recognition task was easier than the category retrieval because both the face and the associated object were presented as retrieval cues.

### Transformation of engram representations revealed by pattern dissimilarity

A common notion in neuroscience is that neurons that encode information also retain the information and are reactivated in the retrieval process (*4*, *13*, *23*, *24*, *62*, *63*). Accordingly, we expected to observe retrieval-related activation within the same voxels that had been activated during encoding. Indeed, voxel patterns repeated from encoding to 30-minute category retrieval within medial temporal and neocortical areas. Yet, the voxel patterns in the hippocampus and anterior cingulate gyrus became dissimilar from the 30-minute to the 24-hour category retrieval for both consciously accessible and inaccessible memories. Hence the retrieval and ensuing re-encoding process at the 30-minute retrieval may have shifted activated voxels within these two key areas of episodic memory, as was observed in the animal model: Neurophysiological studies in animals have shown that hippocampal memory traces can change significantly across time as well as depending on the retrieval demands (*28–31*, *64*, *65*). Lei and colleagues (*31*) recently reported that hippocampal engrams underlying contextual fear memories in mice shift with repeated retrieval. Memory recall can trigger a change in the hippocampal engram leading to the formation of a new, updated hippocampal engram. Ko and colleagues (*66*) also found that the time-dependent transformation of contextual fear memories involves a reorganization of hippocampal engram circuits. For our study, we assume that re-encoding and reconsolidation processes have caused the shift in voxel patterns from the 30-minute to the 24-hour category retrieval. There was no systematic reinstatement or shift of hippocampal engram components from encoding to the 30-minute category retrieval or to the 24-hour category retrieval. However, from encoding to the recognition, where the face and object displayed during encoding were re-presented, a right hippocampal engram component reinstated for sure responses possibly due to visual recollection.

A pattern shift from the 30-minute to the 24-hour category retrieval was not just seen in the right hippocampus but also within bilateral anterior cingulate gyrus. The anterior cingulate gyrus is strongly connected to the hippocampus, heavily involved in early consolidation processes (*20*), shares cell dynamics with the hippocampus (*67*) and even harbors place cells (*68*). The anterior cingulate gyrus is also involved in storing novel information into already established knowledge (*69*). Its prominent role in early memory consolidation, its connectivity with the hippocampus, and its high plasticity may explain why also the anterior cingulate cortex displays a re-encoding and reconsolidation driven retrieval-retrieval dissimilarity.

### Theoretical implications

This research was inspired by experiments in rodents, which suggest that interference-based forgetting is an adaptive form of forgetting, which is reversible and updateable and thus in favor of an inaccessible rather than unavailable engram (*10*). For example, Autore and colleagues (*10*) showed that although natural retroactive interference during learning resulted in forgetting and decreased engram cell reactivation in the retrieval situation, optogenetic stimulation of previously labeled engram cells enabled memory expression. Hence, engram components remained implemented in the brain despite forgetting. In the consciously inaccessible state, as we have shown here, episodic memories can still be reactivated, they can still guide behavior, and they are still subject to active consolidation processes. Hence, they are far from gone. This informs traditional models of memory that distinguish between memory systems based on whether learning and retrieval occur consciously or unconsciously (*33–35*). These models posit that episodic memories are strictly dependent on consciousness, i.e., one needs to have a conscious experience to relive an episode from one’s life. An alternative model suggests that a separation between memory systems based on processing modes is more plausible and is better supported by memory data (*39*). According to this newer model, consciousness modulates memories within each memory system in a quantitative way (i.e., memory strength) but does not qualitatively define memory systems. The current findings provide evidence in favor of this processing-based memory model (*39*) because initially consciously accessible episodic memories can lose conscious accessibility and may regain it when given sufficient retrieval cues. Even those memories that never re-emerge to conscious access remain implemented in the episodic memory network including the hippocampus. These unconscious memories are also subject to active consolidation processes and continue to influence the choice behavior implicitly. A future line of research might examine how long consciously inaccessible memories remain implemented in the brain and whether they can be reinforced brain stimulation to bring them back to conscious access.

## MATERIAL AND METHODS

### EXPERIMENTAL DESIGN

#### Human Participants

Forty healthy, right-handed participants were randomly assigned to two groups. Both groups underwent the same procedures but underwent distinct functional MRI sequences. Twenty participants were examined with a functional MRI sequence with limited FOV covering the medial temporal lobe and adjacent structures (age: 24.30 ± 3.87 (mean ± SD); 10 women). The remaining 20 participants were examined with a whole-brain fMRI sequence (age: 23.8 ± 3.41 (mean ± SD); 11 women). Participants were recruited through advertisements on university- and social-media-platforms. All participants underwent a telephone screening. They reported normal or corrected-to-normal vision, no previous or current neurological or psychiatric diseases, no claustrophobia, and no non-removable metals in or on the body. All participants provided written informed consent for the experiment. Note that the participants’ consent was semi-informed. Participants were left unaware of the 24-hour category retrieval of the face-object combinations learned on the first day. We did not inform them of a 24-hour category retrieval to prevent their active rehearsing of the learning material. Instead, we informed participants that they are reinvited to the second MRI session for them to learn and retrieve another set of face-object combinations. Participants were fully debriefed at the end of the study. Participants were compensated with a reimbursement of 150 Swiss Francs and digital copies of the anatomical scans of their own brains. The study was approved by the ethics committee of the Canton of Berne (ID: 50008).

#### Stimuli

The stimulus set selected for this experiment consisted of 96 combinations of faces and objects. Face stimuli were selected from the Flickr-Faces-HQ Dataset (FFHQ, https://github.com/NVlabs/ffhq-dataset). Selection was restricted to Caucasian faces to reduce bias. Faces were neither standardized nor clipped, and paraphernalia (e.g., a hat or glasses) and the background (e.g., a forest or an office scene) were not removed from the pictures to preserve the stimuli’s naturalistic appearance and to allow for better discriminability of faces. We selected images of everyday objects on white backgrounds from two categories: organic (e.g. an apple, a bird) and inorganic (e.g. a hammer, a couch). The set of 96 faces consisted of 48 male and 48 female faces. Half of the male faces were paired with an organic object and the other half with an inorganic object. The same pairing scheme was applied to the female faces.

##### Stimulus groups

Groups of four stimuli were created because of the constraints of the recognition: During recognition, participants were presented with a face plus two object stimuli (the target object and a foil object). Their task was to recognize the object that was paired with the face during learning. All foil objects had been presented with different face stimuli during learning and were thus familiar. Recognition required participants to remember which familiar object had been presented with the current face during learning (associative recognition). For stimulus preparation, we created groups of two face and two object stimuli. The two face stimuli of a certain group would be presented with the same two objects from their group during the recognition task, one object being the target and the other the foil for the initially presented face and vice versa for the subsequently presented face. To exclude object bias in the groups, we collected data in a pilot study using an online survey. In this survey, 63 students decided which of two objects would fit better to either one of the two face stimuli using a rating scale from 0-100. Based on this survey, we included those groups in the main experiment with the least amount of object bias, i.e., groups that yielded a mean rating score close to 50. Table S19 shows the mean scores for each of the 48 groups used in the main experiment. The 48 groups provided 96 face-object pairs.

##### Randomization

Target and foil objects were randomly assigned to face stimuli within each group. The presentation order and the interval between the two face-object combinations belonging to a group were semi-randomized such that the time interval is maximal between the two combinations that belong to one group. A maximal temporal distance between the two face-object combinations belonging to one group was important for participants’ accuracy on the recognition task because on the recognition task the same two objects from each group were presented with each of the two faces from the group. Maximizing the distance between subsequent presentations increased the likelihood that participants would make their decision of which object was paired with a given face during learning solely based on memory and not based on their previous choice of an object. The order, in which the 96 face-object pairs were presented during learning, was randomized with a windowed shuffle for the 30-minute and the 24-hour category retrieval and for the recognition. The windowed shuffle shuffled 4 list entries before moving the window to the next 4 entries. This ensured that the study-test retention interval was comparable for the 96 memories. Moreover, the shuffling also preserved the distance between the two face-objects pairs from the same group.

#### Experimental tasks

For all tasks, presentations of stimuli were separated by a jittered interstimulus-interval (ISI) of 500 – 2500 ms. A black fixation cross on a white background was presented during the ISI in the center of the screen. Participants were asked to focus on the fixation cross.

##### Associative learning task

First, the face stimulus was presented in the center of the screen for 1000 ms. Participants were instructed to look closely at the face and memorize it. Thereafter, the same face was presented again, this time for 2500 ms paired with an object that was drawn from one of two categories: organic or inorganic. Participants were asked to form an association between the presented face and the object. No instructions were given on how to form an association. Following associative learning, the face was presented in the center of the screen for 2750 ms for participants to indicate by button press whether the object the face had just been paired with was organic or inorganic. This immediate category retrieval served to identify trials with successful learning. Only these trials entered the analysis of the fate of engrams underlying subsequently forgotten memories.

##### Consolidation Task

A 5-minute pause was given following the immediate retrieval to allow for memory consolidation. During this pause we kept participants busy with a simple attention task (odd-even task) to avoid active rehearsal. We presented numbers between 0 and 100 on the screen for participants to indicate whether a number was odd or even. Corrective feedback was provided by a color cue: the number changed to green for correct and to red for incorrect odd-even responses.

##### Category Retrieval Task

Each of the 96 face stimuli was presented in the center of the white screen as a retrieval cue for 1000 ms. Then, the face appeared on the screen again, this time for 2750 ms combined with the written instruction to indicate by button press whether the associated object was organic or inorganic. The index finger was assigned to the response button for inorganic, and the middle finger was assigned to the response button for organic. Each retrieval trial was followed by a confidence rating regarding the just provided retrieval response. Participants were given 2500 ms to provide their confidence response using a 3-point scale (sure, unsure, guess). The unsure option was the conceptually largest category including all responses that were neither completely sure nor pure guesses. We evaluated only the extremes, i.e., the sure and the guess responses because we were interested in contrasting consciously remembered associations (sure responses) with forgotten associations (guess responses). The cued associative 2-alternative forced-choice category retrieval is referred to as category retrieval throughout the manuscript.

##### Recognition Task

Each face was presented with the two objects from its group. The presentation duration was 3500 ms. One of the two objects was the target that had been presented along with the face during learning, and the other was a familiar foil object that was presented with another face during learning. Participants were required to indicate which of the two objects had been presented with the face during learning. The two objects were presented twice (though paired with two distinct faces) during the recognition. Each object was once the target and once the foil. Following each recognition trial, participants gave their confidence assessment on a 3-point scale (sure, unsure, guess). The cued associative 2-alternative forced-choice recognition task is referred to as recognition task throughout the manuscript.

##### Confidence rating

After each retrieval (except the immediate category retrieval) and the recognition task, participants were asked to rate their level of confidence using a 3-point scale. Participants were instructed to rate a retrieval response as “sure” if they could vividly remember the face-object combination. Participants were instructed to rate a retrieval response as “guess” if they had forgotten the face-object combination, forgetting meaning no sense of familiarity, no intuition, no “gut feeling” regarding the face-object combination. Participants were instructed to rate a retrieval as “unsure” if their retrieval experience fell anywhere between “sure” and “guess”. We implemented this broad “unsure” category to sort out all responses with varying grades of conscious access to analyze clearly consciously accessible memories with totally forgotten memories. This procedure ensured that any above-chance performance in the ‘guess’ category was due to unconscious processes alone. We included aggregated type-2 d’ scores in Table S16, which shows that participants used the metacognitive confidence scale as instructed. Type-2 d’ were calculated as described by Fleming and Lau (*70*).

#### Technical Setup

The paradigm was presented with a Hyperion projector from Psychology Software Tools (Sharpsburg, USA) with a resolution of 1920 x 1080p Full HD, a refresh rate of 60 Hz, and dimensions of the projected image being set at 30 cm x 54 cm and a screen diagonal of 62cm. The distance between the participants’ eyes and the projection screen was 1.5 m. The experiment was performed using Presentation® software (Version 23.0, Neurobehavioral Systems, Inc., Berkeley, CA) on a Windows computer. Responses were recorded in the MRI scanner with a hand sleeve with a wrist strap and a Response unit for response registration in the MRI scanner named “Celeritas Fiber Optic Response Unit - Right hand, 5 buttons” from Psychology Software Tools (Sharpsburg, USA). Participants used their index, middle, and ring finger of their right hand to submit responses during the task.

#### Experimental Procedure

The study was conducted on two consecutive days. All participants underwent the same protocol. Whenever participants were required to perform a behavioral task in the scanner, BOLD signal was recorded. The study consisted of 4 behavioral tasks, and 4 BOLD fMRI runs. Between runs, fMRI data acquisition was paused. The first three runs were performed consecutively on the first day and the fourth run was performed on the following day. Between the two MRI sessions there was a 24-hour interval, during which participants were free to follow their usual routines. The experiment would usually start between midday and afternoon. The first MRI session lasted about 1 hour, and the second MRI session lasted about 45 minutes.

##### Day 1

On the first day, participants completed an online training at home to establish familiarity with the behavioral tasks. After arriving at and entering the MRI scanner, a t1-weighted anatomical image was recorded. During the anatomical scan participants rehearsed the behavioral tasks and the button press responses. Then, participants completed the associative learning of 96 face-object pairs, carried out the immediate category retrieval and then took the odd-even task during the 5-minute pause. Finally, they performed the 30-minute category retrieval, which marked the end of the first day of experimentation. To prevent active rehearsal of the learned material, participants were misinformed that they would learn another set of face-object pairs on the next day. Participants were asked not to perform cognitively demanding tasks between the two MRI sessions. At the end of the first experimentation day, participants received an e-mail which included a reminder for the second experimentation day as well as a link to a survey to assess sleep duration and sleep quality (Stanford Sleepiness Scale (SSS) and Stanford Sleepiness Test (SST)). Participants were asked to fill out the survey the next morning.

##### Day 2

When participants arrived at the MR center, they received a debriefing in written form, where we explained that they would not learn a new set of face-object combinations but retrieve the face-object pairs learned on the previous day. When asked about their expectations for the second day, 32.5% of participants reported that they had expected a repeated retrieval of the face-object pairs learned on the previous day. But none of participants reported to have actively rehearsed the face-object pairs. Participants entered the scanner again on the second day and we recorded a whole-brain anatomical t1-weighted scan. Next, participants performed the category retrieval and then the recognition task for the face-object associations that they had learned on the previous day. Each category retrieval trial was immediately followed by its corresponding recognition trial.

#### MRI data acquisition

Both anatomical and functional MRI sequences were recorded on a 7T Siemens Magnetom Terra whole body MRI scanner (Siemens Medical Solutions, Erlangen, Deutschland) with a 32-channel head coil. The anatomical t1-weighted image acquisition followed a magnetization-prepared two rapid gradient echo (MP2RAGE) sequence with a time of repetition (TR) of 6000 ms; echo time (TE) 2.06 ms; flip angle (FA) = 0°; field of view (FOV) = 384 x 384 mm^2^, spatial resolution of 0.63 x 0.63 x 0.63 mm^3^, 256 sagittal slices, matrix points = 610. For the t2*-weighted functional MRI scans, we used two distinct sequences. The small field-of-view (FOV) fMRI sequence covered the bilateral medial temporal lobe with 0.8 mm isotropic voxel size to target hippocampal engram components, and the other fMRI sequence covered the entire brain with a 1.1 mm isotropic voxel size to target neocortical engram components.

##### Small FOV fMRI sequence

Isotropic functional T2* weighted blood oxygen level dependent (BOLD) sensitive multiple interleaved slice acquisition sequence with a TR = 2120 ms; TE = 24 ms; FA = 70°; FOV = 210 x 210 mm^2^, spatial resolution of 0.8 x 0.8 x 0.8 mm^3^, 28 transversal slices, matrix points = 263. We acquired a total of 2100 volumes per participant. Day 1: associative learning, 5-minute pause with odd-even task, and associative category retrieval: 510, 150, 515; day 2: associative category retrieval and associative recognition: 925.

##### Whole-brain fMRI sequence

Isotropic functional T2* weighted blood oxygen level dependent (BOLD) sensitive multiple interleaved slice acquisition sequence with a TR = 2700 ms; TE = 50 ms; FA = 90°; FOV = 210 x 210 mm^2^, spatial resolution of 1.1 x 1.1 x 1.1 mm^3^, 96 transversal slices; matrix points = 191. A total of 1650 volumes were collected per participant. Day 1: associative learning, 5-minute pause with odd-even task, and associative category retrieval: 400, 119, 405, day 2: associative category retrieval and associative recognition: 726.

### QUANTIFICATION AND STATISTICAL ANALYSIS

#### Sample size justification

Studies using 3T MRI scanners usually achieve a sample size of around N = 40 (*71–74*). Like other studies employing ultra-high field fMRI (*75–77*), we implemented a sample size of 20 participants per fMRI sequence, leveraging the superior signal-to-noise ratio of 7T MRI scanners compared to 3T MRI scanners (*21*).. This enabled us to employ the same behavioral paradigm in two distinct participant groups with two distinct fMRI sequences – a high-resolution hippocampal fMRI sequence and a lower resolution whole-brain fMRI sequence.

#### Statistical analysis of behavioral data

Behavioral data were preprocessed and analyzed with custom R scripts using RStudio and the tidyverse package on a MacBook Pro. Face-object pairs were removed from the analysis if participants answered incorrectly during immediate category retrieval, answered incorrectly but with high confidence (sure response) or if they pressed the wrong button. The significance threshold was set to p < 0.05, if not indicated otherwise. The Bayes factor was calculated using the “BayesFactor” R-library which makes use of the function described by Rouder (*78*) and colleagues, with default prior distributions for all parameters.

#### Preprocessing of fMRI data

Whole-brain fMRI sequence: Results included in this manuscript for the whole-brain fMRI sequence come from preprocessing performed using fMRIPrep 20.2.3 (*79*) (RRID: SCR _016216), which is based on Nipype 1.6.1 (*80*, *81*) (RRID _SCR002502).

#### Anatomical data preprocessing

A total of 2 T1-weighted (T1w) images were acquired per participant within the input BIDS dataset. All of them were corrected for intensity non-uniformity (INU) with N4Bias FieldCorrection (*82*), distributed with ANTs 2.3.3 (*83*), (RRID:SCR_004757). The T1w-reference was then skull-stripped with a Nipype implementation of the antsBrainExtraction.sh workflow (from ANTs), using OASIS30ANTs as target template. Brain tissue segmentation of cerebrospinal fluid (CSF), white-matter (WM) and gray-matter (GM) was performed on the brain-extracted T1w using fast (*84*) (FSL 5.0.9, RRID:SCR _002823). A T1w-reference map was computed after registration of 2 T1w images (after INU-correction) using mri_robust _template (*85*) (FreeSurfer 6.0.1). Brain surfaces were reconstructed using recon-all(*86*) (FreeSurfer 6.0.1, RRID:SCR _001847), and the brain mask estimated previously was refined with a custom variation of the method to reconcile ANTs-derived and FreeSurfer-derived segmentations of the cortical gray-matter of Mindboggle (*87*) (RRID:SCR _002438). Volume-based spatial normalization to two standard spaces (MNI152NLin2009cAsym, MNI152NLin6Asym) was performed through nonlinear registration with antsRegistration (ANTs 2.3.3), using brain-extracted versions of both T1w reference and the T1w template. The following templates were selected for spatial normalization: ICBM 152 Nonlinear Asymmetrical template version 2009c (*88*) [RRID:SCR_008796; TemplateFlow ID: MNI152NLin2009cAsym], FSL’s MNI ICBM 152 non-linear 6^th^ Generation Asymmetric Average Brain Stereotaxic Registration Model (*89*) [RRID:SCR _002823; TemplateFlow ID: MNI152NLin6Asym].

#### Functional data preprocessing (whole-brain fMRI sequence)

For each of the 4 BOLD runs acquired per participant (across all tasks and sessions), the following preprocessing was performed. First, a reference volume and its skull-stripped version were generated using a custom methodology of fMRIPrep. Susceptibility distortion correction (SDC) was omitted. The BOLD reference was then co-registered to the T1w reference using bbregister (FreeSurfer) which implements boundary-based registration.(*90*) Co-registration was configured with six degrees of freedom. Head-motion parameters with respect to the BOLD reference (transformation matrices, and six corresponding rotation and translation parameters) are estimated before any spatiotemporal filtering using mcflirt (*91*) (FSL 5.0.9). BOLD runs were slice-time corrected using 3dTshift from AFNI 20160207(*92*) (RRID:SCR _005927). The BOLD time-series were resampled onto the following surfaces (FreeSurfer reconstruction nomenclature): fsnative, fsaverage5. The BOLD time-series (including slice-timing correction when applied) were resampled onto their original, native space by applying the transforms to correct for headmotion. These resampled BOLD time-series will be referred to as preprocessed BOLD in original space, or just preprocessed BOLD. The BOLD time-series were resampled into standard space, generating a preprocessed BOLD run in MNI152NLin2009cAsym space. First, a reference volume and its skull-stripped version were generated using a custom methodology of fMRIPrep. Automatic removal of motion artifacts using independent component analysis (ICA-AROMA (*93*)) was performed on the preprocessed BOLD on MNI space after removal of non-steady state volumes and spatial smoothing with an isotropic Gaussian kernel of 6 mm FWHM (full-width half-maximum). Corresponding “non-aggresively” denoised runs were produced after such smoothing. Additionally, the “aggressive” noise-regressors were collected and placed in the corresponding confounds file. Several confounding time-series were calculated based on the preprocessed BOLD: framewise displacement (FD), DVARS and three region-wise global signals. FD was computed using two formulations following Power (*94*) (absolute sum of relative motions) and Jenkinson (*91*) (relative root mean square displacement between affines). FD and DVARS are calculated for each functional run, both using their implementations in Nipype (following the definitions by Power (*94*) and colleagues). The three global signals are extracted within the CSF, the WM, and the whole-brain masks. Additionally, a set of physiological regressors were extracted to allow for component-based noise correction (CompCor (*95*)). Principal components are estimated after high-pass filtering the preprocessed BOLD time-series (using a discrete cosine filter with 128s cut-off) for the two CompCor variants: temporal (tCompCor) and anatomical (aCompCor). tCompCor components are then calculated from the top 2% variable voxels within the brain mask. For aCompCor, three probabilistic masks (CSF, WM and combined CSF+WM) are generated in anatomical space. The implementation differs from that of Behzadi and colleagues in that insteadof eroding the masks by 2 pixels on BOLD space, the aCompCor masks are subtracted a mask of pixels that likely contain a volume fraction of GM. This mask is obtained by dilating a GM mask extracted from the FreeSurfer’s aseg segmentation, and it ensures components are not extracted from voxels containing a minimal fraction of GM. Finally, these masks are resampled into BOLD space and binarized by thresholding at 0.99 (as in the original implementation). Components are also calculated separately within the WM and CSF masks.

For each CompCor decomposition, the k components with the largest singular values are retained, such that the retained components’ time series are sufficient to explain 50 percent of variance across the nuisance mask (CSF, WM, combined, or temporal). The remaining components are dropped from consideration. The head-motion estimates calculated in the correction step were also placed within the corresponding confounds file. The confound time series derived from head motion estimates and global signals were expanded with the inclusion of temporal derivatives and quadratic terms for each (*96*). Frames that exceeded a threshold of 0.5 mm FD or 1.5 standardised DVARS were annotated as motion outliers. All resamplings can be performed with a single interpolation step by composing all the pertinent transformations (i.e. head-motion transform matrices, susceptibility distortion correction when available, and co-registrations to anatomical and output spaces). Gridded (volumetric) resamplings were performed using antsApplyTransforms (ANTs), configured with Lanczos interpolation to minimize the smoothing effects of other kernels (*97*). Non-gridded (surface) resamplings were performed using mri_vol2surf (FreeSurfer).

Many internal operations of fMRIPrep use Nilearn 0.8.189(*98*) (RRID:SCR_001362), mostly within the functional processing workflow. For more details of the pipeline, see the section corresponding to workflows in fMRIPrep’s documentation.

##### Copyright waiver

The above boilerplate text was automatically generated by fMRIPrep with the express intention that users should copy and paste this text into their manuscripts unchanged. It is released under the CC0 license.

#### Functional data preprocessing (small FOV sequence)

Due to the small FOV of the sequence the use of fMRIprep was not recommended. Preprocessing was done with SPM12 (Welcome Department of Cognitive Neurology, London, UK). The functional volumes were slice-time corrected and realigned to the mean functional volume image. Coregistration to the participants anatomical volume was carried out with a non-skull-stripped structural volume. The latter allowed for a more reliable coregistration because the skull provided for more anatomical information to properly match the small FOV with the anatomical template. Volumes were segmented and spatially normalized to the MNI/ICBM AVG152 Template provided by SPM. Functional volumes were smoothed with a 6 mm (FWHM) isotropic Gaussian Kernel.

#### Freesurfer segmentations

We created subject-specific ROIs for both the functional connectivity analyses as well as the RSA analyses with freesurfer. FMRIprep already applies *recon_all* during its workflow which computes ROIs for the neocortex. Hippocampal ROIs were created with freesurfers *segmentHA_T1* function. The creation of neocortical masks failed for 3 participants from the whole-brain fMRI group. They were thus excluded from the representational similarity analysis which used the masks for region of interest analyses.

#### GLM

In the following, both fMRI sequences (small FOV and whole-brain sequence) were always treated identically unless stated otherwise.

##### First Level

First-level analysis was performed using a general linear model (GLM) that modeled the BOLD-time-series using the sequence of event onsets convolved with the default hemodynamic response function (HRF) provided by SPM12. Events were modeled at a duration of zero. For each participant, 4 timeseries were estimated in the same first-level model. All stimulus presentations were modeled as events, the confidence ratings were modeled as events, and the button presses given during the category retrieval and during recognition were modeled as events. These events were categorized into 5 conditions depending on the responses given during the category retrieval: responses to every trial could be categorized as correct sure responses, incorrect unsure responses, correct unsure responses, incorrect guess responses or correct guess responses. For example, the 30-minute category retrieval responses were used for the categorization of corresponding trials during the encoding time-series, to investigate subsequent memory effects: the presentation of the face stimulus during encoding was modeled as an event and categorized based on the response during the retrieval of said face. Sure but incorrect answers were not estimated because their rate was too low for a robust estimate. Six movement regressors (x, y, z, pitch, yaw, roll) were added to the model for every session as well as a session constant which resulted in 129 estimated beta maps for every participant. Implicit masking with the default SPM threshold of 0.8 resulted in the exclusion of signal from the center of the brain, mainly in the whole-brain fMRI sequence. In both sequences, we used a binary brain mask, generated by SPMs segmentation algorithm, as an explicit mask and set SPM’s masking threshold to [−inf], to include all voxels within the mask in the first level estimation. Note that all events were used for first level specification, including trials that were excluded during the behavioral analysis (e.g. false memories, see above), to allow for a better baseline estimation of the general linear model.

##### Second Level

Second-level, random-effects analyses for group statistics were computed using a full factorial model with two factors. The first factor modeled 5 events of interest: the first presentation of the face stimulus during the learning task, the associative learning event, the presentation of the face stimulus during the 30-minute as well as the 24-hour category retrieval, and the recognition. The second factor modeled the retrieval quality for 3 different levels, correct sure responses, correct guessing responses, and incorrect guessing responses. This resulted in a 5 x 3 second-level full factorial design. The height threshold for the calculation of the t-contrasts was set to p < 0.001, uncorrected for multiple comparisons, with a cluster extent threshold of k = 10 voxels, if not indicated otherwise. Note that for the purpose of preserving transparency about sub-threshold results (*99*), some figures are presented at a height threshold of p = 0.005.

#### Brain-behavior correlations

To isolate brain activity that was associated with task accuracy we used a multiple regression design within SPM. For all the models calculated we included the estimation of an intercept. The generated SPM contrast images were combined with a between-subjects behavioral covariate in a multiple-regression model. The behavioral covariate used for contrasts of guess responses was the z-scored percentage correct of all guess responses (guessing accuracy), for contrasts of sure responses the z-scored absolute number of correct sure responses (sure accuracy), because of a ceiling effect for sure responses, i.e. very few incorrect sure responses. For all second-level models we calculated both positive and negative brain-behavior correlations. The significance level set to p<0.001, uncorrected, with a cluster extent threshold of k = 10 voxels, if not indicated otherwise.

#### Stacked contrasts

To uncover both differences between retrieval quality (sure > guess) as well as retrieval timepoints (24-hour > 30-minute category retrieval), we calculated stacked contrasts by contrasting 4 parameter estimates of the corresponding retrieval timepoints and qualities within SPM12s result interface. The significance threshold was set to p < 0.001, uncorrected, with a cluster extent threshold of k = 10 voxels, if not indicated otherwise.

#### Conjunction analyses

To uncover commonalities of brain-behavior correlations for consciously accessible and for forgotten memories, we calculated conjunction analyses. We used the above generated second level multiple regression models and extracted a binary mask of all the voxels that were significant under the threshold of p<0.001 and an extent of k =10 voxels. Then we took these maps as inclusive masks in another second level model of interest and set the threshold again to p<0.001 with a cluster extent of k = 10. The significant clusters in the resulting t-maps only revealed the overlay of all the voxels that would be significant in both models independently of each other. To uncover commonalities of contrast analyses, we used the contrast maps produced by the second level full factorial model to calculate commonalities between activation patterns using SPM12. The significance threshold was set to p < 0.001, uncorrected, with a cluster extent threshold of k = 10 voxels, if not indicated otherwise.

#### Functional connectivity analyses

To uncover task-related connectivity changes between the hippocampus and the neocortex we calculated general psycho-physiological interactions (gPPI) analyses. We used the conn toolbox (version 22a). Unlike the standard PPI analyses provided by SPM12, the conn toolbox (using a gPPI approach) can model effects of more than two conditions, to allow for interactions between conditions, and to make no assumptions how these conditions are related to the baseline condition. First level specifications were imported from SPM to the conn toolbox. Behavioral measures (sure and guessing accuracy) served as second-level covariates. We defined seed regions within the subdivided left and right hippocampus. More precisely, we used grey matter masks of the hippocampus head, body and tail generated by freesurfer. These seed regions were integrated as participant-specific ROIs. Conn’s integrated setup and denoising pipeline was applied to the data with a bandpass filter between 0.008 – 0.09 and linear detrending but no despiking. We also included the second-level covariates to be able to correlate task related changes in connectivity with behavioral parameters like the accuracy of the guess responses given at the 30-minute and 24-hour category retrieval. We included this in a seed-to-voxel analysis with an underlying multivariate regression model. For conjunctions of multivariate regression functional connectivity analyses, we made use of the shared architecture of the two toolboxes, conn and SPM12. We used the contrast files created by the multivariate regression functional connectivity analyses to calculate commonalities between them using SPM12.

#### Representational similarity analysis

Methodology was modeled after the protocol described by Hebscher and colleagues (*63*). One notable difference was how single-trial beta estimates were created. While Hebscher and colleagues used a least-squares-single (LSS, individual GLMs for each trial), we used a least-squares-all (LSA, one GLM with one condition per trial). We examined reinstatement of memory-specific patterns of neural activity by conducting a series of RSAs on patterns of estimated neural activity within ROIs. RSAs measured encoding-retrieval (or encoding-recognition) similarity by computing the Pearson correlation between patterns for all pairs of trials for each ROI. Pairwise correlation values were Fisher transformed. We then subtracted the mean of all correlations calculated for non-matching trial-pairs from the mean correlation of matching trial-pairs. Matching trials refers to trials containing a presentation of the same face during encoding and retrieval. This metric was z-scored against a null distribution of correlation difference scores, calculated by randomly shuffling trials over 1000 permutations. Group-level statistics were performed on these z-scored pattern similarity scores. Finally, to correct for similarity based on face representation and based on the mere task characteristics or encoding/retrieval effort, we subtracted the similarity values of encoding-retrieval pairs that yielded incorrect guess responses from the similarity values of encoding-retrieval pairs that yielded correct sure responses and correct guess responses. This way, we came up with similarity scores that reflected encoding-retrieval or retrieval-retrieval voxel pattern reinstatements. Pair-wise representational similarity analysis (RSA) was conducted with custom python scripts making use of the software package Nilearn (*100*). Correlation values were calculated with the *pairwise_distances* function of the software package scikit-learn (*101*). To assess the significance of voxel pattern reinstatement, we computed one-sample two-tailed t-tests on the z-scored pattern similarity values for each ROI. Similarity estimates were also correlated across subjects with retrieval accuracy. Results below a threshold of p < 0.05 were considered significant. In addition, we computed the Bayes Factor for all t-tests and across-subjects Pearson correlation using the “pingouin” python package, which computes Wetzels (*102*) Bayes factor. We used default prior distributions for all parameters.

## ACKNOWLEDGEMENTS

This work was supported by Sitem-Insel Support Funds SISF 2019 to K. Henke and by the SNFS Advanced Grant TMAG-1_209374 to K. Henke. The authors thank Mirco Bristle Brogli for his support during data analysis. Calculations were performed on UBELIX (https://www.id.unibe.ch/hpc), the HPC cluster at the University of Bern.

## Author Contributions

Conceptualization: T.W., K.H.; Methodology: T.W., A.F., K.H.; Analysis: T.W., K.Z., F.R., A.F.; Writing - original draft: T.W.; Writing - review & editing: T.W., K.Z., K.H.; Visualization: T.W.; Supervision: K.H.; Project administration: K.H.; Funding acquisition: K.H.

## Data and Code availability

The magnetic resonance imaging data have been deposited at “OpenNeuro” and are publicly available.

- Small FOV fMRI group: https://doi.org/10.18112/openneuro.ds006265.v1.0.1
- Whole-brain fMRI group: https://doi.org/10.18112/openneuro.ds006266.v1.0.1

Custom MATLAB, R, and python analysis scripts as well as the behavioral data are available at https://github.com/TomW92/TOAM-fMRI.

## Declaration of Interests

The authors declare no competing interest.

## Declaration of generative AI and AI-assisted technologies

During the preparation of this work, the author(s) used AI-assisted technologies for the reformatting of tables, to create pictograms for Figure 1, and to generate abbreviations from lists of brain regions and for checking grammar. Thereafter, the author(s) reviewed and edited the content and take(s) full responsibility for the content of the publication.

## SUPPLEMENTARY MATERIAL

Supplementary Material can be found in the pdf accompanying this manuscript. The PDF file includes:

1. Figs S1 to S6
2. Tables S1 to S15

## Notes

https://doi.org/10.18112/openneuro.ds006265.v1.0.2

https://doi.org/10.18112/openneuro.ds006266.v1.0.2

https://github.com/TomW92/TOAM-fMRI

### Competing Interest Statement

The authors have declared no competing interest.

### Summary of Updates

- Added type-2 d' analysis - Revised several paragraphs for clarity - Improved clarity in usage of core concepts - Added Luzius Brogli as author to better reflect contributions to data analysis - Figure 1 has been adapted to show immediate retrieval - Changed layout to reflect journal submission demands

