## Supplementary Figures and Tables for "Neural Traces of Forgotten Memories Persist in Humans and are Behaviorally Relevant"

Tom Willems *et al.*

**This PDF file includes:**

- Figs. S1 to S6
- Tables S1 to S15

### Supplementary Figures

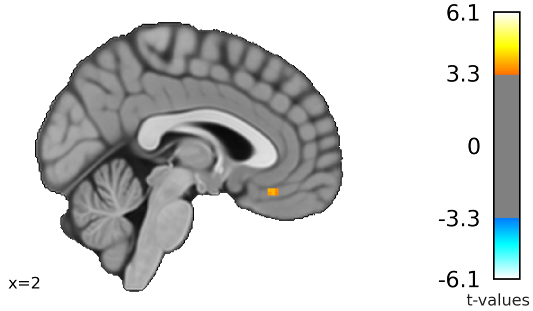


**Figure S1. Medial prefrontal cortex activation during guess responses given at the 24-hour category retrieval predicted the subsequent regain of conscious access to the memories.**

Medial prefrontal component of the episodic retrieval network predicted conscious recognition success. We contrasted the guess responses on the 24-hour category retrieval task that would subsequently yield correct sure responses on the recognition task with those other guess responses given on the 24-hour category task that would subsequently yield correct guess responses on the recognition task. This revealed a cluster in the medial prefrontal cortex (peak at MNI [0, 30, -16], $p_{uncor} < 0.001$, $T(18) = 4.19$). Results presented in this panel were acquired with the whole-brain fMRI sequence. See Table S12 for details.

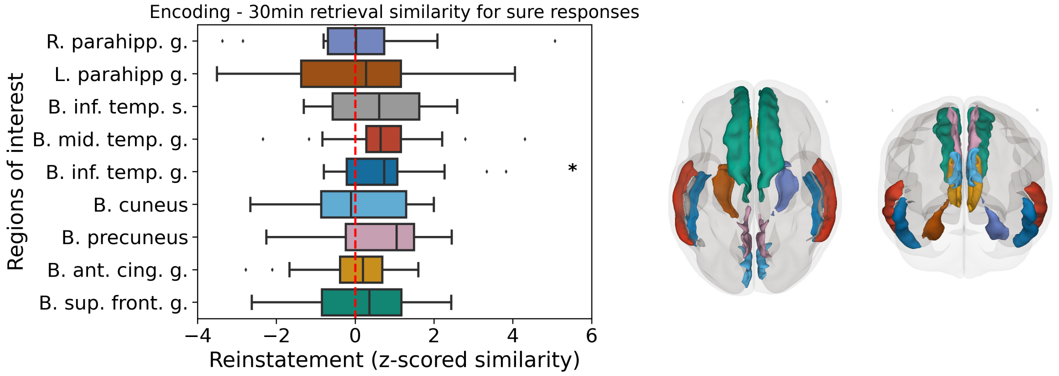


**Figure S2. Similarity of voxel patterns between encoding and the 30-minute category retrieval task was relevant for retrieval accuracy.**

Encoding – retrieval similarity (ERS) analysis for the 30-minute category retrieval yielded significance in the bilateral inferior temporal gyrus for correct sure responses ($t(16) = 2.3$, $p = 0.035$, $B_{10} = 1.9$). Results presented in this panel were acquired with the whole-brain fMRI sequence. *$p < 0.05$, **$p < 0.01$ by Student's t test. Abbreviations: R., right; L., left; B., bilateral; parahipp., parahippocampus; inf., inferior; mid., middle; ant., anterior; cing., cingulate; sup., superior; front., frontal; s., sulcus, g., gyrus.

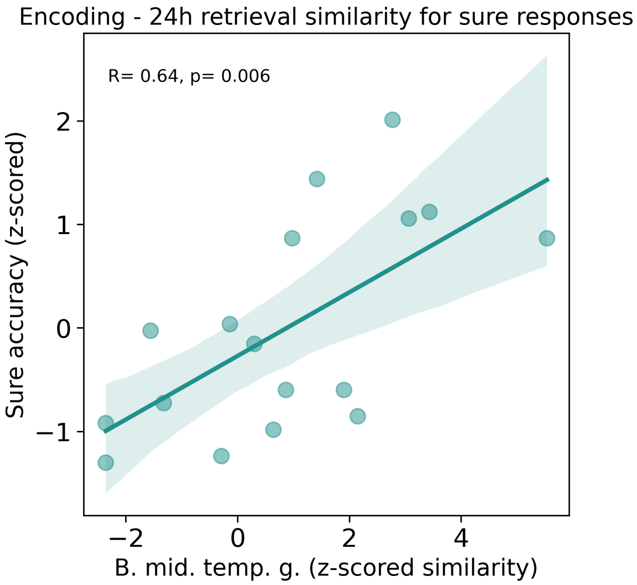


**Figure S3. Similarity of voxel patterns between encoding and the 24-hour category retrieval was relevant for retrieval success.**

Encoding - 24-hour category retrieval similarity in bilateral middle temporal gyrus underlying correct sure responses given on the category retrieval task revealed a non-significant mean value comparison but a significant between-subjects correlation with the accuracy of sure responses given on the 24-hour category task (number of correct sure trials, $p = 0.006$, $R = 0.64$, $B_{10} = 10.4$). Results presented in this panel were acquired with the whole-brain fMRI sequence. Abbreviations: B., bilateral; mid., middle; temp., temporal; g., gyrus.

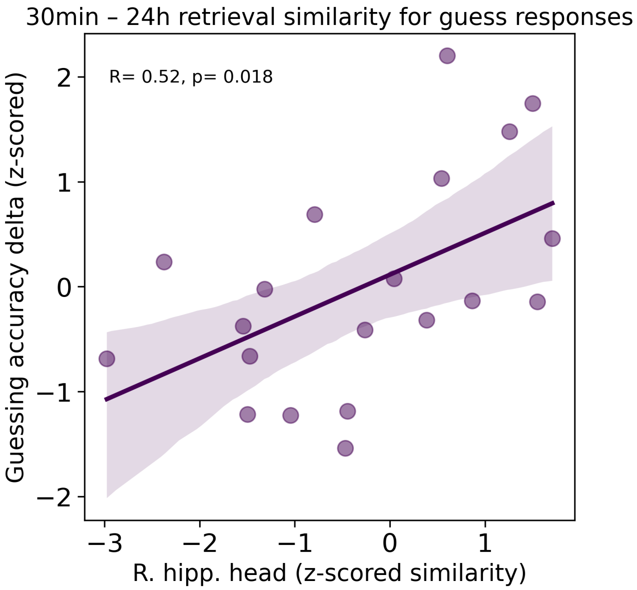


**Figure S4. Similarity of voxel patterns between the 30-minute and the 24-hour category retrieval was relevant for the accuracy of guess responses.**

Retrieval-retrieval similarity of voxel patterns in the right hippocampal head underlying correct guess responses revealed a non-significant mean value comparison but a significant between-subjects correlation with the delta of correct guess responses (i.e., number of correct guess responses at 24 hours minus number of correct guess responses at 30 minutes, $p = 0.018$, $R = 0.52$, $B_{10} = 3.7$). Results presented in this panel were acquired with the small FOV fMRI sequence. Abbreviations: R., right; hipp., hippocampus.

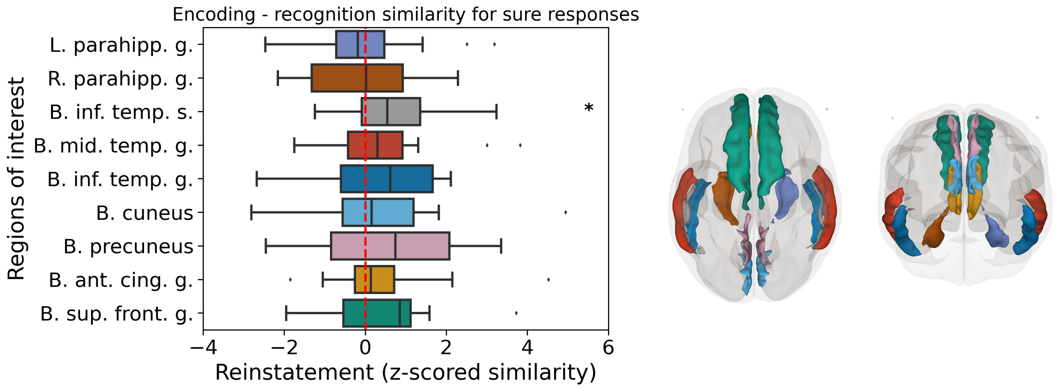


**Figure S5. Similarity of voxel patterns between encoding and recognition at 24 hours was relevant for retrieval success.**

Significant mean value comparison in bilateral inferior temporal sulcus regarding correct sure responses ($p = 0.026$, $t(17) = 2.55$, $B_{10} = 2.9$). Results presented in this panel were acquired with the whole-brain fMRI sequence. *$p < 0.05$, **$p < 0.01$ by Student's t test. Abbreviations: R., right; L., left; B., bilateral; parahipp., parahippocampus; inf., inferior; mid., middle; ant., anterior; cing., cingulate; sup., superior; front., frontal; s., sulcus, g., gyrus.

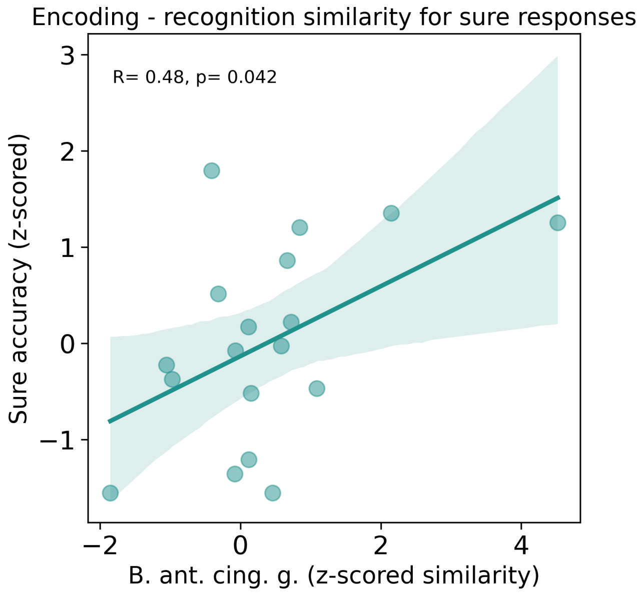


**Figure S6. Similarity of voxel patterns between encoding and recognition at 24 hours was relevant for retrieval success.**

Pattern similarity in bilateral anterior cingulate gyrus (B. ant. cing. g.) underlying correct sure responses revealed a non-significant mean value comparison but a significant between-subjects correlation with the number of correct sure responses given on the recognition task ($p = 0.041$, $R = 0.48$, $B_{10} = 2.0$). Results presented in this panel were acquired with the small FOV fMRI sequence. Abbreviations: B., bilateral; ant., anterior; cing., cingulate; g., gyrus.

### Supplementary Tables

| Cases | Sum of Squares | df | Mean Square | F | p |
|---|---|---|---|---|---|
| Repetition | 0.002 | 1 | 0.002 | 0.041 | 0.840 |
| n | 0.050 | 1 | 0.050 | 1.106 | 0.296 |
| Residuals | 3.464 | 77 | 0.045 | | |

**Table S1. Additional analysis regarding guessing accuracy during the recognition task, related to Figure 2.**
Since objects were repeatedly shown during the recognition task (see STAR Methods), it was theoretically possible to answer recognition trials correctly solely based on previous choices for these objects, while still rating them as guess answers. No participant reported to have done so. Still, we investigated if responses for repeated objects would contribute differently to guessing accuracy than responses for first presentations of the objects (within the recognition task). In addition, we included the number of guessed responses as a covariate, because it varied considerably across participants. The table shows the result of an analysis of covariance (ANCOVA) which shows that it is unlikely that repeated objects (case: repetition) contributed differently to guessed retrieval accuracy, and that number of guessed responses (case: n) was not associated with guessing accuracy across participants.

| Anatomical description | | Cluster | | | | Peak | | | | MNI | | |
|---|---|---|---|---|---|---|---|---|---|---|---|---|
| | | p(FWE-corr) | p(FDR-corr) | k | p(unc) | p(FWE-corr) | p(FDR-corr) | T | p(unc) | x | y | z |
| **R** | Hippocampus | 0.692 | 0.920 | 128 | 0.120 | 0.472 | 0.519 | 4.05 | 0.000 | 30 | -32 | -9 |
| **L** | Posterior cingulate gyrus | 0.998 | 0.920 | 13 | 0.630 | 0.999 | 0.963 | 3.25 | 0.001 | -9 | -52 | 4 |

**Table S2. Common hippocampal and neocortical areas mediated correct sure and correct guess responses at the 30-minute category retrieval, related to Figure 3A.** This table shows the results of a conjunction analysis for guessed correct > guessed incorrect and sure correct > guessed incorrect trials during 30-min category retrieval. This analysis displays the shared brain activation patterns for guessed correct and sure correct trials.

*Notes*. Results from small FOV fMRI sequence. Threshold set to $p < .001$, $k = 10$ voxels, uncorrected. MNI, Montreal Neurological Institute; FWE-corr, family wise error corrected; FDR-corr, false discovery rate corrected; unc, uncorrected; g, gyrus; k, cluster size in number of voxels; L, left hemisphere; R, right hemisphere; B, bilateral.

| Anatomical description | | Cluster | | | | Peak | | | | MNI | | |
|---|---|---|---|---|---|---|---|---|---|---|---|---|
| | | p(FWE-corr) | p(FDR-corr) | k | p(unc) | p(FWE-corr) | p(FDR-corr) | T | p(unc) | x | y | z |
| **L** | Lingual gyrus | 0.996 | 0.920 | 18 | 0.563 | 0.999 | 0.985 | 3.28 | 0.001 | -16 | -54 | 1 |
| **L** | Posterior cingulate gyrus | 1.000 | 0.920 | 2 | 0.876 | 1.000 | 0.985 | 3.16 | 0.001 | -11 | -45 | -1 |
| **L** | Lingual gyrus | 1.000 | 0.920 | 1 | 0.920 | 1.000 | 0.985 | 3.13 | 0.001 | -17 | -55 | 2 |

**Table S3. Common neocortical areas mediated correct sure and correct guess responses at the 24-hour category retrieval, related to Figure 3B.**
This table shows the results of a conjunction analysis for guessed correct > guessed incorrect and sure correct > guessed incorrect trials during 24-hour category retrieval. This analysis displays the shared brain activation patterns for guessed correct and sure correct trials.

*Notes*. Results from small FOV fMRI sequence. Threshold set to $p < .001$, $k = 10$ voxels. MNI, Montreal Neurological Institute; FWE-corr, family wise error corrected; FDR-corr, false discovery rate corrected; unc, uncorrected; g, gyrus; k, cluster size in number of voxels; L, left hemisphere; R, right hemisphere; B, bilateral.

| Anatomical description | | Cluster | | | | Peak | | | | MNI | | |
|---|---|---|---|---|---|---|---|---|---|---|---|---|
| | | p(FWE-corr) | p(FDR-corr) | k | p(unc) | p(FWE-corr) | p(FDR-corr) | T | p(unc) | x | y | z |
| **R** | Hippocampus | 0.0188 | 0.0734 | 431 | 0.0012 | 0.1792 | 0.4407 | 6.3158 | 0.0000 | 27 | -8 | -25 |
| **L** | Ventral diencephalon | 0.1765 | 0.2571 | 227 | 0.0127 | 0.2362 | 0.4407 | 6.1201 | 0.0000 | -16 | -14 | -13 |
| **R** | Middle temporal g. | 0.4410 | 0.3754 | 147 | 0.0382 | 0.2712 | 0.4407 | 6.0196 | 0.0000 | 48 | -49 | 14 |
| **R** | Ventral diencephalon | 0.3373 | 0.3186 | 171 | 0.0270 | 0.2881 | 0.4407 | 5.9748 | 0.0000 | 1 | -18 | -11 |
| **L** | Inferior temporal g. | 0.9870 | 0.8894 | 35 | 0.2852 | 0.3332 | 0.4407 | 5.8651 | 0.0000 | -56 | -10 | -35 |
| **L** | Inferior temporal g. | 0.3259 | 0.3186 | 174 | 0.0259 | 0.7606 | 0.8631 | 5.0893 | 0.0000 | -44 | -27 | -22 |
| **R** | Middle temporal g. | 0.9551 | 0.8894 | 50 | 0.2037 | 0.8645 | 0.8631 | 4.8843 | 0.0001 | 59 | -9 | -32 |
| **R** | Precunues | 0.9457 | 0.8894 | 53 | 0.1912 | 0.8899 | 0.8631 | 4.8239 | 0.0001 | 4 | -64 | 18 |
| **L** | Posterior cingulate g. | 0.9842 | 0.8894 | 37 | 0.2720 | 0.9171 | 0.8631 | 4.7497 | 0.0001 | -5 | -51 | 12 |
| **R** | Middle temporal g. | 0.9809 | 0.8894 | 39 | 0.2597 | 0.9184 | 0.8631 | 4.7459 | 0.0001 | 64 | -56 | 8 |
| **L** | Middle temporal g. | 0.9870 | 0.8894 | 35 | 0.2852 | 0.9478 | 0.8631 | 4.6455 | 0.0001 | -52 | -14 | -25 |
| **R** | Hippocampus | 0.9551 | 0.8894 | 50 | 0.2037 | 0.9524 | 0.8631 | 4.6264 | 0.0001 | 16 | -39 | 6 |
| **L** | Middle temporal g. | 0.1806 | 0.2571 | 225 | 0.0131 | 0.9535 | 0.8631 | 4.6219 | 0.0001 | -54 | -19 | -15 |
| | Brain Stem | 0.5792 | 0.4788 | 121 | 0.0568 | 0.9714 | 0.8631 | 4.5312 | 0.0001 | 0 | -24 | -20 |
| **L** | Middle temporal g. | 0.6320 | 0.4839 | 112 | 0.0656 | 0.9716 | 0.8631 | 4.5300 | 0.0001 | -68 | -28 | -16 |
| **L** | Ventral diencephalon | 0.9968 | 0.8894 | 24 | 0.3770 | 0.9991 | 0.9201 | 4.1062 | 0.0003 | -11 | -18 | -16 |
| **L** | Middle temporal g. | 0.9895 | 0.8894 | 33 | 0.2993 | 0.9994 | 0.9201 | 4.0704 | 0.0004 | -58 | -29 | -10 |

**Table S4. Activity in hippocampus and episodic memory network underlying correct guess responses correlated with the guessing accuracy on the category retrieval task at 30 minutes, related to Figure 3C.**
This table shows the results of a brain-behavior correlation for correct > incorrect guess responses correlated with guessed retrieval performance at 30-minute retrieval. This analysis displays brain activation during 30-min retrieval that was modulated by guessed retrieval performance at 30-min retrieval.

*Notes*. Results from small FOV fMRI sequence. Threshold set to $p < .001$, $k = 10$ voxels. MNI, Montreal Neurological Institute; FWE-corr, family wise error corrected; FDR-corr, false discovery rate corrected; unc, uncorrected; g, gyrus; k, cluster size in number of voxels; L, left hemisphere; R, right hemisphere; B, bilateral; ROI, region of interest.

| Anatomical description | | Cluster | | | | Peak | | | | MNI | | |
|---|---|---|---|---|---|---|---|---|---|---|---|---|
| | | p(FWE-corr) | p(FDR-corr) | k | p(unc) | p(FWE-corr) | p(FDR-corr) | T | p(unc) | x | y | z |
| **R** | Middle temporal gyrus | 0.528 | 0.544 | 148 | 0.060 | 0.320 | 0.775 | 5.77 | 0.000 | 60 | -24 | -11 |
| **R** | Hippocampus | 0.814 | 0.544 | 89 | 0.135 | 0.421 | 0.775 | 5.55 | 0.000 | 31 | -36 | -6 |
| **R** | Hippocampus | 0.931 | 0.563 | 60 | 0.214 | 0.890 | 0.775 | 4.70 | 0.000 | 28 | -10 | -28 |
| **R** | Middle temporal gyrus | 0.997 | 0.715 | 20 | 0.477 | 0.908 | 0.775 | 4.65 | 0.000 | 68 | -22 | -18 |
| **L** | Temporal pole | 0.851 | 0.544 | 81 | 0.152 | 0.941 | 0.775 | 4.55 | 0.000 | -36 | 4 | -29 |
| **R** | Middle temporal gyrus | 0.995 | 0.715 | 25 | 0.423 | 0.955 | 0.775 | 4.49 | 0.000 | 56 | -16 | -15 |
| **R** | Middle temporal gyrus | 0.974 | 0.612 | 43 | 0.291 | 0.968 | 0.775 | 4.43 | 0.000 | 61 | -16 | -16 |
| | Brain stem | 0.833 | 0.544 | 85 | 0.143 | 0.980 | 0.775 | 4.35 | 0.000 | -2 | -36 | -10 |
| **L** | Hippocampus | 0.968 | 0.612 | 46 | 0.275 | 0.991 | 0.775 | 4.24 | 0.000 | -17 | -40 | 3 |
| **L** | Middle temporal gyrus | 0.896 | 0.544 | 70 | 0.181 | 0.992 | 0.775 | 4.21 | 0.000 | -52 | -46 | 4 |
| | Brain stem | 0.997 | 0.715 | 22 | 0.454 | 0.996 | 0.775 | 4.14 | 0.000 | 9 | -19 | -27 |

**Table S5. Hippocampal and middle temporal gyrus activity underlying correct guess responses correlated with the guessing accuracy on the category retrieval task at 24 hours, related to Figure 3C.**
This table shows the results of a brain-behavior correlation for correct > incorrect guess responses that correlated with guessed retrieval performance at 24-hour retrieval. This analysis displays brain activation during 24-hour guessed retrieval that was modulated by guessed retrieval performance.

*Notes*. Results from small FOV fMRI sequence. Clusters identified solely within white matter regions have been excluded. Threshold set to $p < .001$, $k = 10$ voxels. MNI, Montreal Neurological Institute; FWE-corr, family wise error corrected; FDR-corr, false discovery rate corrected; unc, uncorrected; g, gyrus; k, cluster size in number of voxels; L, left hemisphere; R, right hemisphere; B, bilateral; ROI, region of interest.

| Anatomical description | | Cluster | | | | Peak | | | | MNI | | |
|---|---|---|---|---|---|---|---|---|---|---|---|---|
| | | p(FWE-corr) | p(FDR-corr) | k | p(unc) | p(FWE-corr) | p(FDR-corr) | T | p(unc) | x | y | z |
| **B** | Anterior cingulate g. & medial superior frontal g. | 0.000 | 0.000 | 86 | 0.000 | 0.042 | 0.363 | 7.81 | 0.000 | 12 | 42 | -4 |
| | | | | | | 1.000 | 0.918 | 4.69 | 0.000 | 12 | 42 | 10 |
| **R** | Precuneus | 0.704 | 0.083 | 13 | 0.008 | 0.400 | 0.474 | 6.61 | 0.000 | 14 | -48 | 52 |
| **L** | Medial superior frontal g. | 0.929 | 0.132 | 10 | 0.017 | 0.958 | 0.474 | 6.12 | 0.000 | -6 | 34 | 30 |
| **L** | Middle cingulate gyrus | 0.000 | 0.000 | 60 | 0.000 | 0.986 | 0.479 | 5.93 | 0.000 | -6 | -22 | 34 |
| | | | | | | 1.000 | 0.853 | 5.10 | 0.000 | 6 | -22 | 42 |
| | | | | | | 1.000 | 0.897 | 4.89 | 0.000 | -12 | -32 | 42 |
| **R** | Anterior cingulate gyrus | 0.704 | 0.083 | 13 | 0.008 | 0.995 | 0.541 | 5.79 | 0.000 | 8 | 34 | 6 |
| **R** | Cuneus | 0.704 | 0.083 | 13 | 0.008 | 0.998 | 0.589 | 5.67 | 0.000 | 4 | -82 | 6 |
| **R** | Precuneus | 0.262 | 0.052 | 19 | 0.002 | 1.000 | 0.827 | 5.40 | 0.000 | 12 | -60 | 26 |
| | Cerebellar vermal lobules | 0.448 | 0.063 | 16 | 0.004 | 1.000 | 0.853 | 5.27 | 0.000 | 4 | -46 | -18 |
| | Brain stem | 0.377 | 0.057 | 17 | 0.003 | 1.000 | 0.853 | 5.25 | 0.000 | -2 | -32 | -6 |
| **L** | Precuneus | 0.377 | 0.057 | 17 | 0.003 | 1.000 | 0.853 | 5.23 | 0.000 | -8 | -60 | 34 |
| | | | | | | 1.000 | 0.897 | 4.75 | 0.000 | -14 | -64 | 42 |
| **L** | Precuneus | 0.614 | 0.077 | 14 | 0.006 | 1.000 | 0.853 | 5.13 | 0.000 | -20 | -58 | 24 |
| **L** | Middle cingulate gyrus | 0.528 | 0.067 | 15 | 0.005 | 1.000 | 0.853 | 5.12 | 0.000 | -8 | 0 | 36 |
| **L** | Lingual gyrus | 0.148 | 0.045 | 22 | 0.001 | 1.000 | 0.853 | 5.03 | 0.000 | -14 | -54 | 0 |
| | | | | | | 1.000 | 0.953 | 4.09 | 0.000 | -4 | -52 | 4 |
| **L** | Precuneus | 0.262 | 0.052 | 19 | 0.002 | 1.000 | 0.853 | 5.01 | 0.000 | -12 | -54 | 20 |
| **L** | Hippocampus | 0.868 | 0.127 | 11 | 0.013 | 1.000 | 0.853 | 5.00 | 0.000 | -34 | -28 | -10 |
| | | | | | | 1.000 | 0.953 | 4.11 | 0.000 | -28 | -32 | -6 |
| **L** | Posterior cingulate gyrus | 0.315 | 0.057 | 18 | 0.002 | 1.000 | 0.875 | 4.97 | 0.000 | -2 | -44 | 24 |
| **L** | Cuneus | 0.929 | 0.132 | 10 | 0.017 | 1.000 | 0.897 | 4.81 | 0.000 | 0 | -84 | 36 |
| **L** | Anterior cingulate gyrus | 0.262 | 0.052 | 19 | 0.002 | 1.000 | 0.897 | 4.75 | 0.000 | -12 | 42 | 8 |
| **L** | Thalamus | 0.929 | 0.132 | 10 | 0.017 | 1.000 | 0.897 | 4.75 | 0.000 | -18 | -24 | 10 |
| **R** | Lingual gyrus | 0.016 | 0.007 | 34 | 0.000 | 1.000 | 0.918 | 4.59 | 0.000 | 12 | -52 | 2 |
| **R** | Posterior cingulate gyrus | 0.929 | 0.132 | 10 | 0.017 | 1.000 | 0.918 | 4.50 | 0.000 | 4 | -38 | 28 |
| **R** | Pallidum | 0.377 | 0.057 | 17 | 0.003 | 1.000 | 0.918 | 4.47 | 0.000 | 14 | -8 | -4 |
| **L** | Anterior insula | 0.929 | 0.132 | 10 | 0.017 | 1.000 | 0.918 | 4.44 | 0.000 | -30 | 20 | -10 |
| **L** | Precuneus | 0.448 | 0.063 | 16 | 0.004 | 1.000 | 0.918 | 4.43 | 0.000 | -4 | -46 | 46 |
| | | | | | | 1.000 | 0.953 | 4.17 | 0.000 | -8 | -44 | 38 |
| **R** | Superior frontal gyrus | 0.929 | 0.132 | 10 | 0.017 | 1.000 | 0.945 | 4.25 | 0.000 | 16 | 52 | 34 |

**Table S6. Hippocampal functional connectivity during learning correlated with the 30-minute guessing accuracy for forgotten associations, related to Figure 4A.**
This table shows the results of a correlative gPPI analysis with the seed region in the right hippocampus head: learning trials that would yield correct > incorrect guess responses during the 30-minute retrieval (subsequent memory analysis) revealed that connectivity estimates correlated between-subjects with guessing accuracy. This analysis displays right hippocampus head functional connectivity during learning that was modulated by subsequent guessing accuracy.

*Notes*. Results from whole-brain fMRI sequence. Clusters identified solely within white matter regions have been excluded. Threshold set to $p < .001$, $k = 10$ voxels. MNI, Montreal Neurological Institute; FWE-corr, family wise error corrected; FDR-corr, false discovery rate corrected; unc, uncorrected; g, gyrus; gPPI, general psychophysiological interaction; k, cluster size in number of voxels; L, left hemisphere; R, right hemisphere; B, bilateral.

| Anatomical description | | Cluster | | | | Peak | | | | | MNI | | |
|---|---|---|---|---|---|---|---|---|---|---|---|---|---|
| | | p(FWE-corr) | p(FDR-corr) | k | p(unc) | p(FWE-corr) | p(FDR-corr) | T | Z | p(unc) | x | y | z |
| **L** | Temporal pole | 0.920 | 0.138 | 10 | 0.016 | 0.014 | 0.327 | 8.41 | 5.29 | 0.000 | -48 | 18 | -22 |
| **L** | Middle frontal gyrus | 0.920 | 0.138 | 10 | 0.016 | 0.336 | 0.811 | 6.70 | 4.69 | 0.000 | -26 | 42 | 16 |
| **L** | Inferior temporal gyrus | 0.201 | 0.044 | 20 | 0.001 | 0.393 | 0.811 | 6.62 | 4.65 | 0.000 | -48 | -40 | -32 |
| **L** | Caudate | 0.042 | 0.020 | 28 | 0.000 | 0.396 | 0.811 | 6.62 | 4.65 | 0.000 | -16 | -8 | 18 |
| **R** | Medial superior frontal gyrus | 0.000 | 0.000 | 144 | 0.000 | 0.820 | 0.813 | 6.25 | 4.50 | 0.000 | 12 | 46 | 28 |
| | | | | | | 1.000 | 0.957 | 5.41 | 4.11 | 0.000 | 8 | 48 | 36 |
| | | | | | | 1.000 | 0.957 | 5.30 | 4.06 | 0.000 | 6 | 36 | 20 |
| **R** | Medial superior frontal g. | 0.425 | 0.081 | 16 | 0.004 | 0.992 | 0.957 | 5.87 | 4.33 | 0.000 | -4 | 56 | 18 |
| **R** | Inferior temporal gyrus | 0.355 | 0.074 | 17 | 0.003 | 0.995 | 0.957 | 5.80 | 4.30 | 0.000 | 54 | -4 | -40 |
| **R** | Middle temporal gyrus | 0.092 | 0.034 | 24 | 0.001 | 0.997 | 0.957 | 5.77 | 4.29 | 0.000 | 62 | -42 | 2 |
| **R** | Posterior insula | 0.504 | 0.082 | 15 | 0.005 | 1.000 | 0.957 | 5.53 | 4.18 | 0.000 | 38 | -12 | 16 |
| **L** | Anterior cingulate gyrus | 0.504 | 0.082 | 15 | 0.005 | 1.000 | 0.957 | 5.53 | 4.17 | 0.000 | -4 | 36 | 8 |
| **R** | Middle temporal gyrus | 0.111 | 0.034 | 23 | 0.001 | 1.000 | 0.957 | 5.44 | 4.13 | 0.000 | 66 | -8 | -22 |
| **L** | Precentral gyrus | 0.591 | 0.100 | 14 | 0.006 | 1.000 | 0.957 | 5.43 | 4.13 | 0.000 | -38 | -12 | 62 |
| | | | | | | 1.000 | 0.957 | 4.33 | 3.54 | 0.000 | 18 | 50 | -6 |
| **L** | Posterior orbital gyrus | 0.682 | 0.112 | 13 | 0.007 | 1.000 | 0.957 | 5.18 | 4.00 | 0.000 | -30 | 28 | -20 |
| **R** | Posterior orbital gyrus | 0.042 | 0.020 | 28 | 0.000 | 1.000 | 0.957 | 5.13 | 3.98 | 0.000 | 32 | 26 | -22 |
| **L** | Temporal pole | 0.029 | 0.020 | 30 | 0.000 | 1.000 | 0.957 | 5.03 | 3.92 | 0.000 | -42 | 16 | -40 |
| **R** | Inferior frontal gyrus | 0.000 | 0.000 | 80 | 0.000 | 1.000 | 0.957 | 4.97 | 3.89 | 0.000 | 50 | 26 | 10 |
| | | | | | | 1.000 | 0.957 | 4.95 | 3.88 | 0.000 | 40 | 30 | 0 |
| | | | | | | 1.000 | 0.970 | 3.95 | 3.31 | 0.000 | 50 | 28 | 0 |
| **L** | Anterior cingulate gyrus | 0.201 | 0.044 | 20 | 0.001 | 1.000 | 0.957 | 4.97 | 3.89 | 0.000 | -8 | 20 | -10 |
| **R** | Middle temporal gyrus | 0.772 | 0.124 | 12 | 0.010 | 1.000 | 0.957 | 4.94 | 3.88 | 0.000 | 52 | -60 | 6 |
| **L** | Medial orbital gyrus | 0.136 | 0.034 | 22 | 0.001 | 1.000 | 0.957 | 4.86 | 3.83 | 0.000 | -18 | 26 | -26 |
| **L** | Medial precentral gyrus | 0.920 | 0.138 | 10 | 0.016 | 1.000 | 0.957 | 4.85 | 3.83 | 0.000 | -10 | -18 | 52 |
| **L** | Precuneus | 0.920 | 0.138 | 10 | 0.016 | 1.000 | 0.957 | 4.62 | 3.70 | 0.000 | -14 | -66 | 36 |
| **L** | Postcentral gyrus | 0.854 | 0.129 | 11 | 0.012 | 1.000 | 0.957 | 4.56 | 3.67 | 0.000 | -58 | -16 | 48 |
| **R** | Supplementary motor cortex | 0.504 | 0.082 | 15 | 0.005 | 1.000 | 0.957 | 4.55 | 3.66 | 0.000 | 12 | 4 | 52 |
| **L** | Frontal operculum | 0.355 | 0.074 | 17 | 0.003 | 1.000 | 0.957 | 4.52 | 3.64 | 0.000 | -42 | 12 | 0 |
| | | | | | | 1.000 | 0.970 | 3.85 | 3.24 | 0.001 | -36 | 12 | -6 |
| **R** | Caudate | 0.854 | 0.129 | 11 | 0.012 | 1.000 | 0.957 | 4.50 | 3.64 | 0.000 | 8 | 6 | 12 |
| **R** | Superior frontal gyrus | 0.425 | 0.081 | 16 | 0.004 | 1.000 | 0.957 | 4.47 | 3.62 | 0.000 | 24 | 12 | 62 |
| **L** | Superior temporal gyrus | 0.682 | 0.112 | 13 | 0.007 | 1.000 | 0.957 | 4.35 | 3.55 | 0.000 | -68 | -26 | -2 |
| **R** | Posterior orbital gyrus | 0.920 | 0.138 | 10 | 0.016 | 1.000 | 0.957 | 4.31 | 3.53 | 0.000 | 32 | 26 | -12 |
| **R** | Middle frontal gyrus | 0.772 | 0.124 | 12 | 0.010 | 1.000 | 0.957 | 4.21 | 3.47 | 0.000 | 36 | 0 | 56 |
| **R** | Supplementary motor cortex | 0.920 | 0.138 | 10 | 0.016 | 1.000 | 0.959 | 4.18 | 3.45 | 0.000 | -6 | 20 | 62 |
| **R** | Anterior cingulate gyrus | 0.920 | 0.138 | 10 | 0.016 | 1.000 | 0.970 | 4.08 | 3.39 | 0.000 | 6 | 44 | 12 |
| **L** | Postcentral gyrus | 0.854 | 0.129 | 11 | 0.012 | 1.000 | 0.970 | 4.08 | 3.39 | 0.000 | -62 | -14 | 24 |

**Table S7. Hippocampal functional connectivity at the 30-minute retrieval correlated with the 30-minute guessing accuracy, related to Figure 4B.**
This table shows the results of a correlative gPPI analysis with the seed region in the right hippocampus head: correct > incorrect guess responses correlated with 30-min guessing accuracy. This analysis displays right hippocampus head connectivity during correct > incorrect guess responses given at the 30-minute category retrieval correlated with guessing accuracy at 30 minutes.

*Notes*. Results from whole-brain fMRI sequence. Clusters identified solely within white matter regions have been excluded. Threshold set to $p < .001$, $k = 10$ voxels. MNI, Montreal Neurological Institute; FWE-corr, family wise error corrected; FDR-corr, false discovery rate corrected; unc, uncorrected; g, gyrus; gPPI, general psychophysiological interaction; k, cluster size in number of voxels; L, left hemisphere; R, right hemisphere; B, bilateral;

| Anatomical description | | Cluster | | | | Peak | | | | MNI | | |
|---|---|---|---|---|---|---|---|---|---|---|---|---|
| | | p(FWE-corr) | p(FDR-corr) | k | p(unc) | p(FWE-corr) | p(FDR-corr) | T | p(unc) | x | y | z |
| **L** | Superior parietal and supramarginal g. | 0.0005 | 0.0003 | 188 | 0.0000 | 0.4554 | 0.9960 | 4.6550 | 0.0000 | -28 | -44 | 42 |
| **L** | Central operculum | 0.0000 | 0.0000 | 293 | 0.0000 | 0.6020 | 0.9960 | 4.5396 | 0.0000 | -38 | -2 | 22 |
| **R** | Superior frontal g. | 0.6136 | 0.1233 | 49 | 0.0054 | 0.9579 | 0.9960 | 4.1787 | 0.0000 | 24 | 54 | 4 |
| **L** | Superior frontal g. | 0.8631 | 0.2035 | 39 | 0.0112 | 0.9686 | 0.9960 | 4.1517 | 0.0000 | -16 | 12 | 50 |
| **R** | Occipital fusiform & lingual g. | 0.3603 | 0.0784 | 60 | 0.0025 | 0.9958 | 0.9960 | 4.0079 | 0.0000 | 26 | -74 | -8 |
| **R** | Cerebellum exterior | 0.7949 | 0.1697 | 42 | 0.0090 | 0.9960 | 0.9960 | 4.0057 | 0.0000 | 22 | -76 | -26 |
| **R** | Middle frontal g. | 0.0001 | 0.0001 | 223 | 0.0000 | 0.9966 | 0.9960 | 3.9962 | 0.0001 | 32 | 12 | 36 |
| **R** | Precuneus | 0.3797 | 0.0784 | 59 | 0.0027 | 0.9987 | 0.9960 | 3.9445 | 0.0001 | 10 | -58 | 54 |
| **L** | Middle & superior temporal g. | 0.1971 | 0.0541 | 71 | 0.0012 | 0.9998 | 0.9960 | 3.8583 | 0.0001 | -46 | -52 | 14 |
| **L** | Cerebellum exterior | 0.2907 | 0.0695 | 64 | 0.0019 | 0.9999 | 0.9960 | 3.8437 | 0.0001 | -42 | -52 | -30 |
| **L** | Lingual g. and calcarine cortex | 0.1185 | 0.0388 | 80 | 0.0007 | 0.9999 | 0.9960 | 3.8399 | 0.0001 | 18 | -60 | 2 |
| **L** | Anterior insula | 0.1761 | 0.0530 | 73 | 0.0011 | 0.9999 | 0.9960 | 3.8366 | 0.0001 | -34 | 14 | -2 |
| **R** | Occipital pole | 0.8833 | 0.2035 | 38 | 0.0121 | 0.9999 | 0.9960 | 3.8243 | 0.0001 | 16 | -96 | 18 |
| **L** | Angular g. | 0.6926 | 0.1453 | 46 | 0.0067 | 1.0000 | 0.9960 | 3.7882 | 0.0001 | -38 | -70 | 28 |
| **B** | Lingual g. | 0.9473 | 0.2457 | 34 | 0.0166 | 1.0000 | 0.9960 | 3.6585 | 0.0002 | -2 | -80 | -8 |
| **R** | Precentral g. | 0.9589 | 0.2457 | 33 | 0.0180 | 1.0000 | 0.9960 | 3.5930 | 0.0002 | 20 | -22 | 50 |
| **L** | Middle frontal g. | 0.3069 | 0.0695 | 63 | 0.0021 | 1.0000 | 0.9960 | 3.5871 | 0.0002 | -30 | 14 | 30 |
| **R** | Middle frontal g. | 0.5621 | 0.1130 | 51 | 0.0047 | 1.0000 | 0.9960 | 3.5019 | 0.0003 | 30 | 46 | 8 |

**Table S8. Overnight consolidation brought a neocorticalization for sure responses.**
The table shows the results of a stacked contrast (24-hour category retrieval: correct sure > correct guess) > (30-minute category retrieval: correct sure > correct guess responses). This analysis displays overnight changes in activation differences between sure and guessed answers.

*Notes*. Results from whole-brain fMRI sequence. Clusters identified solely within white matter regions have been excluded. Threshold set to $p < .001$, $k = 10$ voxels. MNI, Montreal Neurological Institute; FWE-corr, family wise error corrected; FDR-corr, false discovery rate corrected; unc, uncorrected; g, gyrus; k, cluster size in number of voxels; L, left hemisphere; R, right hemisphere; B, bilateral.

| Anatomical description | Cluster | | | | Peak | | | | MNI | | |
|---|---|---|---|---|---|---|---|---|---|---|---|
| | p(FWE-corr) | p(FDR-corr) | k | p(unc) | p(FWE-corr) | p(FDR-corr) | T | p(unc) | x | y | z |
| **L** Hippocampus | 0.175 | 0.176 | 323 | 0.020 | 0.074 | 0.134 | 4.62 | 0.000 | -27 | -16 | -14 |
| **R** Hippocampus | 0.780 | 0.460 | 106 | 0.154 | 0.629 | 0.445 | 3.92 | 0.000 | 39 | -22 | -12 |
| **L** Inferior temporal gyrus | 0.995 | 0.810 | 20 | 0.540 | 0.914 | 0.486 | 3.63 | 0.000 | -48 | -24 | -26 |

**Table S9. Overnight consolidation brought a deeper and broader hippocampal implementation for guess responses.**
This table shows the results of a contrast analysis of 24-hour category retrieval correctly guessed responses > 30-minutes correctly guessed responses. This analysis displays the overnight changes in brain activation for the guessed trials.

*Notes*. Results from small FOV fMRI sequence. Clusters identified solely within white matter regions have been excluded. Threshold set to p < .001, k = 10 voxels. MNI, Montreal Neurological Institute; FWE-corr, family wise error corrected; FDR-corr, false discovery rate corrected; unc, uncorrected; g, gyrus; k, cluster size in number of voxels; L, left hemisphere; R, right hemisphere; B, bilateral.

| Anatomical description | Cluster | | | | Peak | | | | MNI | | |
|---|---|---|---|---|---|---|---|---|---|---|---|
| | p(FWE-corr) | p(FDR-corr) | k | p(unc) | p(FWE-corr) | p(FDR-corr) | T | p(unc) | x | y | z |
| **L** Entorhinal area & temporal pole | 0.266 | 0.189 | 266 | 0.032 | 0.052 | 0.025 | 4.72 | 0.000 | -33.2 | 1.2 | -26.4 |
| **L** Middle temporal gyrus | 0.716 | 0.384 | 122 | 0.128 | 0.399 | 0.155 | 4.11 | 0.000 | -60.4 | -1.2 | -24.8 |

**Table S10. Overnight consolidation brought a deeper and broader hippocampal implementation for guess responses: no hippocampal clusters for 30-minute > 24-hour retrieval contrast.**
This table shows the results of a contrast analysis of 30-minute category retrieval correctly guessed responses > 24-hours correctly guessed responses. This analysis displays the overnight differences between the two category retrievals.

*Notes*. Results from small FOV fMRI sequence. Threshold set to p < .001, k = 10 voxels. MNI, Montreal Neurological Institute; FWE-corr, family wise error corrected; FDR-corr, false discovery rate corrected; unc, uncorrected; g, gyrus; k, cluster size in number of voxels; L, left hemisphere; R, right hemisphere; B, bilateral.

| Anatomical description | | Cluster | | | | Peak | | | | MNI | | |
|---|---|---|---|---|---|---|---|---|---|---|---|---|
| | | p(FWE-corr) | p(FDR-corr) | k | p(unc) | p(FWE-corr) | p(FDR-corr) | T | p(unc) | x | y | z |
| **R** | Entorhinal area | 0.291 | 0.160 | 22 | 0.008 | 0.781 | 0.999 | 5.770 | 0.000 | 24 | 0 | -16 |
| **L** | Hippocampus and thalamus | 0.011 | 0.016 | 49 | 0.000 | 0.944 | 0.999 | 5.370 | 0.000 | -26 | -30 | -4 |
| | | | | | | 1.000 | 0.999 | 3.800 | 0.001 | -24 | -20 | -2 |
| **L** | Putamen | 0.000 | 0.001 | 83 | 0.000 | 0.986 | 0.999 | 5.120 | 0.000 | -28 | -16 | -8 |
| | | | | | | 0.998 | 0.999 | 4.890 | 0.000 | -30 | -8 | -6 |
| | | | | | | 1.000 | 0.999 | 4.470 | 0.000 | -18 | -8 | -10 |
| **L** | Superior temporal gyrus | 0.074 | 0.072 | 33 | 0.002 | 0.992 | 0.999 | 5.040 | 0.000 | -60 | -14 | -4 |
| **R** | Hippocampus | 0.580 | 0.242 | 16 | 0.020 | 0.996 | 0.999 | 4.950 | 0.000 | 36 | -22 | -12 |
| **R** | Superior temporal gyrus | 0.291 | 0.160 | 22 | 0.008 | 0.998 | 0.999 | 4.890 | 0.000 | 52 | 2 | -18 |
| **R** | Thalamus proper | 0.522 | 0.229 | 17 | 0.017 | 1.000 | 0.999 | 4.710 | 0.000 | 12 | -18 | 4 |
| **R** | Inferior temporal gyrus | 0.820 | 0.398 | 12 | 0.040 | 1.000 | 0.999 | 4.130 | 0.000 | 46 | -16 | -32 |
| **R** | Transverse temporal gyrus | 0.872 | 0.441 | 11 | 0.048 | 1.000 | 0.999 | 4.060 | 0.000 | 38 | -26 | 14 |

**Table S11. Overnight functional connectivity increases improved the guessing accuracy at the 24-hours category retrieval, related to Figure 5.**
This table shows the results of a a gPPI analysis with the seed region in the right hippocampus head using the small FOV fMRI sequence to examine overnight changes in the functional connectivity within the medial temporal lobe underlying correct guess responses. We contrasted the functional connectivity underlying the correct guess responses given at the 24-hour category retrieval with the functional connectivity underlying correct guess responses given at the 30-minute category retrieval and correlated the results with the 24-hours guessing accuracy.

*Notes*. Results from small FOV fMRI sequence. Threshold set to $p < .005$, $k = 20$ voxels. MNI, Montreal Neurological Institute; FWE-corr, family wise error corrected; FDR-corr, false discovery rate corrected; unc, uncorrected; g, gyrus; gPPI, general psychophysiological interaction; k, cluster size in number of voxels; L, left hemisphere; R, right hemisphere; B, bilateral.

| Anatomical description | | Cluster | | | | Peak | | | | MNI | | |
|---|---|---|---|---|---|---|---|---|---|---|---|---|
| | | p(FWE-corr) | p(FDR-corr) | k | p(unc) | p(FWE-corr) | p(FDR-corr) | T | p(unc) | x | y | z |
| **R** | Medial prefrontal cortex | 0.998 | 0.778 | 10 | 0.312 | 1.000 | 0.989 | 4.19 | 0.000 | 0 | 30 | -16 |

**Table S12. Medial prefrontal cortex activation during guess responses given at the 24-hour category retrieval predicted the subsequent regain of conscious access to the memories, related to Figure S1A.**
This table shows the results of a contrast analysis for the guess responses on the 24-hour category retrieval task that would subsequently yield correct sure responses (i.e., conscious access) on the recognition task with those other guess responses given on the 24-hour category retrieval task that would yield correct guess responses (i.e., no conscious access) on the recognition task.

*Notes*. Results from whole-brain fMRI sequence. Threshold set to $p < .001$, $k = 10$ voxels. MNI, Montreal Neurological Institute; FWE-corr, family wise error corrected; FDR-corr, false discovery rate corrected; unc, uncorrected; g, gyrus; k, cluster size in number of voxels; L, left hemisphere; R, right hemisphere; B, bilateral.

| Anatomical description | | Cluster | | | | Peak | | | | MNI | | |
|---|---|---|---|---|---|---|---|---|---|---|---|---|
| | | p(FWE-corr) | p(FDR-corr) | k | p(unc) | p(FWE-corr) | p(FDR-corr) | T | p(unc) | x | y | z |
| **L** | Superior parietal lobule | 0.005 | 0.004 | 43 | 0.000 | 0.017 | 0.129 | 6.61 | 0.000 | -44 | -40 | 54 |
| | | | | | | 1.000 | 0.427 | 4.70 | 0.000 | -40 | -40 | 46 |
| | | | | | | 1.000 | 0.776 | 3.81 | 0.000 | -42 | -48 | 52 |
| **R** | Superior parietal lobule | 0.000 | 0.000 | 116 | 0.000 | 0.022 | 0.129 | 6.52 | 0.000 | 24 | -68 | 54 |
| | | | | | | 0.226 | 0.227 | 5.74 | 0.000 | 26 | -62 | 48 |
| | | | | | | 0.999 | 0.421 | 4.79 | 0.000 | 36 | -70 | 52 |
| **R** | Precuneus | 0.076 | 0.023 | 27 | 0.001 | 0.085 | 0.199 | 6.07 | 0.000 | 8 | -68 | 62 |
| | | | | | | 0.861 | 0.281 | 5.29 | 0.000 | 0 | -66 | 58 |
| **L** | Superior occipital gyrus | 0.064 | 0.022 | 28 | 0.000 | 0.111 | 0.207 | 5.98 | 0.000 | -12 | -90 | 40 |
| | | | | | | 1.000 | 0.574 | 4.36 | 0.000 | -20 | -84 | 40 |
| **L** | Superior occipital gyrus | 0.593 | 0.103 | 15 | 0.006 | 0.144 | 0.220 | 5.89 | 0.000 | -16 | -84 | 48 |
| | | | | | | 1.000 | 0.557 | 4.41 | 0.000 | -22 | -76 | 48 |
| **R** | Superior occipital gyrus | 0.949 | 0.182 | 10 | 0.021 | 0.169 | 0.220 | 5.84 | 0.000 | 16 | -86 | 30 |
| **R** | Precentral gyrus | 0.513 | 0.100 | 16 | 0.005 | 0.231 | 0.227 | 5.74 | 0.000 | 44 | 6 | 34 |
| **L** | Fusiform gyrus & hippocampus | 0.000 | 0.000 | 63 | 0.000 | 0.335 | 0.245 | 5.61 | 0.000 | -30 | -32 | -20 |
| | | | | | | 0.348 | 0.245 | 5.60 | 0.000 | -26 | -32 | -6 |
| | | | | | | 1.000 | 0.599 | 4.32 | 0.000 | -34 | -36 | -6 |
| **L** | Middle temporal gyrus | 0.949 | 0.182 | 10 | 0.021 | 0.337 | 0.245 | 5.61 | 0.000 | -56 | -72 | 6 |
| **R** | Superior frontal gyrus | 0.091 | 0.023 | 26 | 0.001 | 0.398 | 0.255 | 5.55 | 0.000 | 26 | -2 | 60 |
| **R** | Superior occipital gyrus | 0.016 | 0.006 | 36 | 0.000 | 0.727 | 0.281 | 5.35 | 0.000 | 26 | -82 | 44 |
| | | | | | | 1.000 | 0.504 | 4.48 | 0.000 | 34 | -80 | 46 |
| **R** | Precuneus | 0.593 | 0.103 | 15 | 0.006 | 0.826 | 0.281 | 5.31 | 0.000 | 16 | -60 | 26 |
| **L** | Middle temporal gyrus | 0.677 | 0.114 | 14 | 0.008 | 0.831 | 0.281 | 5.31 | 0.000 | -68 | -48 | 4 |
| **L** | Supplementary motor cortex | 0.014 | 0.006 | 37 | 0.000 | 0.875 | 0.281 | 5.27 | 0.000 | -12 | 6 | 56 |
| **L** | Middle frontal gyrus | 0.016 | 0.006 | 36 | 0.000 | 0.972 | 0.374 | 5.04 | 0.000 | -40 | 30 | 20 |
| **L** | Lingual gyrus | 0.759 | 0.125 | 13 | 0.010 | 0.972 | 0.374 | 5.04 | 0.000 | 2 | -84 | 0 |
| | | | | | | 1.000 | 0.826 | 3.66 | 0.000 | 8 | -88 | -6 |
| **R** | Angular gyrus | 0.091 | 0.023 | 26 | 0.001 | 0.978 | 0.384 | 5.02 | 0.000 | 50 | -52 | 34 |
| **L** | Superior parietal lobule | 0.109 | 0.023 | 25 | 0.001 | 0.995 | 0.417 | 4.87 | 0.000 | -32 | -70 | 34 |
| **L** | Calcarine cortex | 0.901 | 0.165 | 11 | 0.016 | 0.996 | 0.417 | 4.86 | 0.000 | -6 | -82 | 4 |
| **R** | Precuneus | 0.187 | 0.034 | 22 | 0.001 | 0.997 | 0.417 | 4.84 | 0.000 | 6 | -74 | 52 |
| **R** | Middle frontal gyrus | 0.156 | 0.030 | 23 | 0.001 | 0.997 | 0.417 | 4.83 | 0.000 | 50 | 42 | 16 |
| | | | | | | 1.000 | 0.718 | 4.06 | 0.000 | 42 | 40 | 16 |
| **L** | Inferior occipital gyrus | 0.109 | 0.023 | 25 | 0.001 | 0.998 | 0.417 | 4.81 | 0.000 | -46 | -82 | 4 |
| | | | | | | 1.000 | 0.815 | 3.69 | 0.000 | -42 | -88 | 8 |
| **R** | Middle frontal gyrus | 0.006 | 0.004 | 42 | 0.000 | 0.999 | 0.423 | 4.76 | 0.000 | 30 | 34 | 40 |
| | | | | | | 1.000 | 0.814 | 3.69 | 0.000 | 34 | 44 | 38 |
| | | | | | | 1.000 | 0.889 | 3.54 | 0.001 | 34 | 36 | 48 |
| **R** | Hippocampus | 0.513 | 0.100 | 16 | 0.005 | 0.999 | 0.427 | 4.74 | 0.000 | 16 | -34 | 10 |
| **L** | Ventral diencephalon | 0.593 | 0.103 | 15 | 0.006 | 1.000 | 0.427 | 4.68 | 0.000 | -2 | -24 | -10 |
| **R** | Inferior temporal gyrus | 0.156 | 0.030 | 23 | 0.001 | 1.000 | 0.427 | 4.66 | 0.000 | 36 | -10 | -46 |
| **L** | Supramarginal gyrus | 0.901 | 0.165 | 11 | 0.016 | 1.000 | 0.427 | 4.66 | 0.000 | -68 | -36 | 22 |
| **L** | Middle cingulate gyrus | 0.836 | 0.154 | 12 | 0.013 | 1.000 | 0.429 | 4.64 | 0.000 | -16 | -16 | 42 |
| **R** | Precentral gyrus | 0.316 | 0.059 | 19 | 0.003 | 1.000 | 0.432 | 4.63 | 0.000 | 50 | 6 | 48 |
| **L** | Superior parietal lobule | 0.949 | 0.182 | 10 | 0.021 | 1.000 | 0.432 | 4.61 | 0.000 | -22 | -62 | 42 |
| **L** | Cerebellum exterior | 0.949 | 0.182 | 10 | 0.021 | 1.000 | 0.563 | 4.38 | 0.000 | -50 | -56 | -28 |
| **L** | Angular gyrus | 0.677 | 0.114 | 14 | 0.008 | 1.000 | 0.563 | 4.37 | 0.000 | -46 | -52 | 54 |
| | | | | | | 1.000 | 0.627 | 4.23 | 0.000 | -38 | -48 | 46 |
| **L** | Superior temporal gyrus | 0.901 | 0.165 | 11 | 0.016 | 1.000 | 0.563 | 4.37 | 0.000 | -66 | -54 | 18 |
| **L** | Postcentral gyrus | 0.677 | 0.114 | 14 | 0.008 | 1.000 | 0.574 | 4.35 | 0.000 | -50 | -24 | 52 |
| | | | | | | 1.000 | 0.754 | 3.94 | 0.000 | -58 | -20 | 52 |
| **R** | Precentral gyrus | 0.901 | 0.165 | 11 | 0.016 | 1.000 | 0.644 | 4.21 | 0.000 | 44 | -4 | 60 |
| | | | | | | 1.000 | 0.776 | 3.88 | 0.000 | 38 | 0 | 64 |
| **R** | Middle occipital gyrus | 0.901 | 0.165 | 11 | 0.016 | 1.000 | 0.708 | 4.09 | 0.000 | 40 | -80 | 40 |
| | | | | | | 1.000 | 0.755 | 3.93 | 0.000 | 40 | -70 | 42 |
| **L** | Precentral gyrus | 0.949 | 0.182 | 10 | 0.021 | 1.000 | 0.770 | 3.90 | 0.000 | -44 | 4 | 34 |
| **R** | Precentral gyrus | 0.901 | 0.165 | 11 | 0.016 | 1.000 | 0.776 | 3.82 | 0.000 | 32 | -14 | 62 |

**Table S13. Hippocampal connectivity to fronto-parietal network mediated recognition success for guess responses.**
This table shows the results of a correlative gPPI analysis with the seed region in the right hippocampus head: correct versus incorrect guess responses during the recognition task correlated with guessed recognition accuracy. This analysis displays right hippocampus head connectivity underlying guess responses that was modulated by number of correct sure responses.

*Notes*. Results from the whole-brain fMRI sequence. Clusters identified solely within white matter regions have been excluded. Threshold set to $p < .001$, $k = 10$ voxels. MNI, Montreal Neurological Institute; FWE-corr, family wise error corrected; FDR-corr, false discovery rate corrected; unc, uncorrected; g, gyrus; gPPI, general psychophysiological interaction; k, cluster size in number of voxels; L, left hemisphere; R, right hemisphere; B, bilateral.

| | Anatomical description | Cluster | | | | Peak | | | | MNI | | |
|---|---|---|---|---|---|---|---|---|---|---|---|---|
| | | p(FWE-corr) | p(FDR-corr) | k | p(unc) | p(FWE-corr) | p(FDR-corr) | T | p(unc) | x | y | z |
| **L** | Lingual gyrus | 0.000 | 0.000 | 99 | 0.000 | 0.008 | 0.129 | 6.88 | 0.000 | -12 | -68 | -8 |
| | | | | | | 1.000 | 0.576 | 4.72 | 0.000 | -22 | -64 | 2 |
| | | | | | | 1.000 | 0.821 | 4.22 | 0.000 | -20 | -72 | -2 |
| **R** | Lingual gyrus | 0.023 | 0.004 | 34 | 0.000 | 0.102 | 0.204 | 6.01 | 0.000 | 6 | -58 | 4 |
| | | | | | | 1.000 | 0.954 | 3.47 | 0.001 | 10 | -50 | 4 |
| **R** | Calcarine cortex | 0.000 | 0.000 | 256 | 0.000 | 0.104 | 0.204 | 6.00 | 0.000 | 6 | -80 | 4 |
| | | | | | | 0.891 | 0.576 | 5.24 | 0.000 | -4 | -86 | -2 |
| | | | | | | 0.972 | 0.576 | 5.04 | 0.000 | 14 | -82 | 8 |
| **R** | Thalamus proper | 0.000 | 0.000 | 74 | 0.000 | 0.105 | 0.204 | 6.00 | 0.000 | 18 | -16 | 2 |
| | | | | | | 0.999 | 0.576 | 4.78 | 0.000 | 12 | -10 | -4 |
| **R** | Occipital pole | 0.006 | 0.001 | 42 | 0.000 | 0.132 | 0.204 | 5.92 | 0.000 | 16 | -98 | 12 |
| | | | | | | 1.000 | 0.821 | 4.22 | 0.000 | 18 | -96 | 20 |
| **L** | Transverse temporal gyrus | 0.016 | 0.003 | 36 | 0.000 | 0.149 | 0.204 | 5.88 | 0.000 | -40 | -32 | 12 |
| | | | | | | 1.000 | 0.707 | 4.47 | 0.000 | -46 | -28 | 6 |
| **L** | Calcarine cortex | 0.003 | 0.001 | 47 | 0.000 | 0.474 | 0.455 | 5.49 | 0.000 | -12 | -80 | 10 |
| | | | | | | 0.999 | 0.576 | 4.78 | 0.000 | -12 | -74 | 2 |
| **L** | Middle and superior frontal g. | 0.000 | 0.000 | 102 | 0.000 | 0.904 | 0.576 | 5.22 | 0.000 | -20 | 52 | 4 |
| | | | | | | 1.000 | 0.797 | 4.35 | 0.000 | -26 | 56 | 12 |
| | | | | | | 1.000 | 0.822 | 4.12 | 0.000 | -30 | 58 | 4 |
| **L** | Planum polare | 0.836 | 0.148 | 12 | 0.013 | 0.965 | 0.576 | 5.07 | 0.000 | -44 | -2 | -12 |
| **R** | Inferior temporal gyrus | 0.759 | 0.129 | 13 | 0.010 | 0.967 | 0.576 | 5.07 | 0.000 | 62 | -16 | -38 |
| **R** | Cuneus | 0.949 | 0.205 | 10 | 0.021 | 0.973 | 0.576 | 5.04 | 0.000 | 4 | -70 | 26 |
| **R** | Middle temporal gyrus | 0.006 | 0.001 | 42 | 0.000 | 0.975 | 0.576 | 5.03 | 0.000 | 62 | -44 | -8 |
| **R** | Cuneus | 0.091 | 0.015 | 26 | 0.001 | 0.978 | 0.576 | 5.02 | 0.000 | 6 | -92 | 24 |
| | | | | | | 1.000 | 0.841 | 4.08 | 0.000 | 8 | -94 | 16 |
| **L** | Transverse temporal gyrus | 0.836 | 0.148 | 12 | 0.013 | 0.985 | 0.576 | 4.98 | 0.000 | -44 | -16 | 4 |
| **R** | Inferior occipital gyrus | 0.223 | 0.030 | 21 | 0.002 | 0.987 | 0.576 | 4.96 | 0.000 | 44 | -78 | 2 |
| **L** | Inferior occipital gyrus | 0.759 | 0.129 | 13 | 0.010 | 0.992 | 0.576 | 4.92 | 0.000 | -44 | -84 | -4 |
| **L** | Middle frontal gyrus | 0.000 | 0.000 | 83 | 0.000 | 0.994 | 0.576 | 4.90 | 0.000 | -32 | 20 | 42 |
| | | | | | | 1.000 | 0.865 | 3.82 | 0.000 | -30 | 16 | 50 |
| **R** | Temporal pole | 0.000 | 0.000 | 74 | 0.000 | 0.996 | 0.576 | 4.87 | 0.000 | 50 | 8 | -38 |
| | | | | | | 1.000 | 0.755 | 4.42 | 0.000 | 48 | 8 | -46 |
| **L** | Putamen | 0.513 | 0.076 | 16 | 0.005 | 0.997 | 0.576 | 4.83 | 0.000 | -34 | -16 | -4 |
| **L** | Anterior cingulate gyrus | 0.001 | 0.000 | 55 | 0.000 | 0.999 | 0.576 | 4.77 | 0.000 | -6 | 44 | 6 |
| | | | | | | 1.000 | 0.821 | 4.30 | 0.000 | -8 | 52 | 10 |
| **R** | Middle temporal gyrus | 0.949 | 0.205 | 10 | 0.021 | 0.999 | 0.576 | 4.77 | 0.000 | 68 | -46 | 8 |
| **R** | Thalamus proper | 0.091 | 0.015 | 26 | 0.001 | 1.000 | 0.579 | 4.68 | 0.000 | 8 | -12 | 6 |
| **L** | Posterior insula | 0.677 | 0.115 | 14 | 0.008 | 1.000 | 0.653 | 4.56 | 0.000 | -38 | -18 | 22 |
| **R** | Precuneus | 0.001 | 0.000 | 55 | 0.000 | 1.000 | 0.684 | 4.52 | 0.000 | 6 | -54 | 38 |
| | | | | | | 1.000 | 0.904 | 3.75 | 0.000 | 10 | -50 | 52 |
| **R** | Central operculum | 0.223 | 0.030 | 21 | 0.002 | 1.000 | 0.706 | 4.49 | 0.000 | 54 | 8 | 0 |
| **R** | Medial frontal gyrus | 0.156 | 0.024 | 23 | 0.001 | 1.000 | 0.755 | 4.41 | 0.000 | 2 | 30 | -18 |
| **L** | Occipital pole | 0.014 | 0.003 | 37 | 0.000 | 1.000 | 0.755 | 4.41 | 0.000 | -12 | -96 | 18 |
| | | | | | | 1.000 | 0.836 | 4.10 | 0.000 | -4 | -90 | 16 |
| | | | | | | 1.000 | 0.841 | 3.99 | 0.000 | -8 | -94 | 10 |
| **R** | Medial superior frontal gyrus | 0.513 | 0.076 | 16 | 0.005 | 1.000 | 0.797 | 4.34 | 0.000 | 8 | 60 | 26 |
| **R** | Anterior cingulate gyrus | 0.949 | 0.205 | 10 | 0.021 | 1.000 | 0.822 | 4.14 | 0.000 | 0 | 34 | -2 |
| **R** | Gyrus rectus | 0.901 | 0.178 | 11 | 0.016 | 1.000 | 0.841 | 4.07 | 0.000 | 6 | 54 | -24 |
| **L** | Thalamus proper | 0.949 | 0.205 | 10 | 0.021 | 1.000 | 0.841 | 4.03 | 0.000 | -16 | -24 | -2 |
| **L** | Middle temporal gyrus | 0.374 | 0.054 | 18 | 0.003 | 1.000 | 0.841 | 4.02 | 0.000 | -64 | -36 | -4 |
| **R** | Anterior cingulate gyrus | 0.901 | 0.178 | 11 | 0.016 | 1.000 | 0.855 | 3.92 | 0.000 | 6 | 46 | 4 |
| **L** | Middle cingulate gyrus | 0.223 | 0.030 | 21 | 0.002 | 1.000 | 0.855 | 3.92 | 0.000 | -2 | 18 | 34 |
| | | | | | | 1.000 | 0.860 | 3.88 | 0.000 | -12 | 12 | 32 |
| | | | | | | 1.000 | 0.904 | 3.69 | 0.000 | -4 | 8 | 36 |

**Table S14. Hippocampal connectivity to fronto-parietal network mediated recognition success for sure responses.**
This table shows the results of a correlative gPPI analysis with the seed region in the right hippocampus head: correct sure versus incorrect guess responses during the recognition task correlated with the number of sure recognized trials. This analysis displays right hippocampus head connectivity underlying sure responses that was modulated by number of correct sure responses.

*Notes*. Results from the whole-brain fMRI sequence. Clusters identified solely within white matter regions have been excluded. Threshold set to $p < .001$, $k = 10$ voxels. MNI, Montreal Neurological Institute; FWE-corr, family wise error corrected; FDR-corr, false discovery rate corrected; unc, uncorrected; g, gyrus; gPPI, general psychophysiological interaction; k, cluster size in number of voxels; L, left hemisphere; R, right hemisphere; B, bilateral.

| Group index | Mean group ratings | SEM of mean group ratings |
|---|---|---|
| 1 | 47.9 | 0.8 |
| 2 | 62.9 | 1.5 |
| 3 | 57.7 | 1.9 |
| 4 | 64.6 | 1.9 |
| 5 | 52.1 | 3.7 |
| 6 | 52.1 | 1.5 |
| 7 | 42.4 | 1.4 |
| 8 | 49.1 | 1.2 |
| 9 | 52.3 | 1.5 |
| 10 | 56.2 | 1.5 |
| 11 | 50.1 | 1.1 |
| 12 | 49.2 | 0.9 |
| 13 | 48.2 | 1.4 |
| 14 | 63.3 | 0.8 |
| 15 | 55.5 | 1.2 |
| 16 | 48.9 | 1.4 |
| 17 | 45.9 | 1.2 |
| 18 | 55.8 | 1.2 |
| 19 | 46.4 | 0.8 |
| 20 | 47.5 | 1.2 |
| 21 | 50.6 | 1.1 |
| 22 | 49.6 | 1.0 |
| 23 | 48.7 | 0.9 |
| 24 | 63.4 | 0.8 |
| 25 | 59.5 | 2.3 |
| 26 | 48.9 | 1.4 |
| 27 | 55.8 | 1.2 |
| 28 | 46.8 | 1.2 |
| 29 | 56.2 | 0.9 |
| 30 | 49.7 | 0.9 |
| 31 | 55.5 | 1.2 |
| 32 | 63.5 | 2.0 |
| 33 | 58.6 | 0.9 |
| 34 | 62.7 | 2.8 |
| 35 | 73.5 | 2.0 |
| 36 | 60.3 | 1.0 |
| 37 | 46.2 | 1.2 |
| 38 | 69.9 | 2.1 |
| 39 | 46.8 | 1.2 |
| 40 | 55.0 | 3.1 |
| 41 | 54.4 | 1.5 |
| 42 | 46.7 | 3.5 |
| 43 | 54.5 | 2.6 |
| 44 | 47.4 | 1.2 |
| 45 | 50.6 | 1.2 |
| 46 | 44.5 | 0.9 |
| 47 | 47.1 | 1.4 |
| 48 | 40.7 | 2.2 |

**Table S15. Online-survey results.**
This table shows the result of an online-survey in which participants (N=63) rated the fit of 2 objects to 2 faces on a scale from 0 to 100. Stimuli-fit was rated equally for a group if the mean group rating was close to 50. We included groups with the least amount of bias in the final stimulus set presented in this table.

*Notes*. SEM: standard error of the mean.

| Task | Rating | HR | FAR | Type 2 d’, M | Type 2 d’, SD |
|---|---|---|---|---|---|
| 30-min retrieval | guess | 0.492 | 0.508 | -0.047 | 0.585 |
| 30-min retrieval | unsure | 0.578 | 0.422 | 0.419 | 0.621 |
| 30-min retrieval | sure | 0.836 | 0.164 | 2.228 | 1.097 |
| 24-hour retrieval | guess | 0.494 | 0.506 | -0.032 | 0.371 |
| 24-hour retrieval | unsure | 0.545 | 0.455 | 0.235 | 0.484 |
| 24-hour retrieval | sure | 0.854 | 0.146 | 2.465 | 1.22 |
| Recognition | guess | 0.555 | 0.445 | 0.308 | 0.757 |
| Recognition | unsure | 0.749 | 0.251 | 1.474 | 0.88 |
| Recognition | sure | 0.945 | 0.055 | 3.549 | 0.948 |

**Table S16. Metacognitive sensitivity (type-2 d’) across retrieval and recognition tasks as a function of confidence rating**.
Data are collapsed across participants for 30-min cateogory retrieval, 24-hour category retrieval, and the recognition task. For each task, mean hit rate (HR) and mean false alarm rate (FAR) across participants are reported for ‘guess’, ‘unsure’ and ‘sure’ confidence categories, alongside the corresponding type-2 d’ measure of metacognitive sensitivity. M and SD are used to represent mean and standard deviation, respectively.
